# Harmonizing brain rhythms: cortex-wide neuronal dynamics underpin quasi-periodic patterns in resting-state fMRI

**DOI:** 10.64898/2026.03.26.713939

**Authors:** Francesca Mandino, Xilin Shen, Corey Horien, Xenophon Papademetris, Stephen M. Strittmatter, Shella D. Keilholz, Nan Xu, Evelyn M.R. Lake

## Abstract

Functional magnetic resonance imaging (fMRI) captures whole-brain activity fluctuations non-invasively in humans and animals. Beyond task/stimuli-locked responses, fMRI measures large-scale patterned activity during rest. Among these, quasi-periodic patterns (QPPs) represent recurring waves of activity that unfold over seconds and exhibit consistent spatiotemporal characteristics. Notably, certain fMRI-QPPs are well-preserved across species and altered in various neuropsychiatric and neurodegenerative diseases. Yet, our collective understanding of their neural underpinnings is limited given the indirect nature of blood-oxygen-level dependent (BOLD) fMRI signals. Simultaneous measures of local field potentials have provided some affirmation that fMRI-QPPs have neural origins, but these point-measurements are limited to a handful of sites. Here, we use a unique multimodal implementation of simultaneous wide-field calcium (WF-Ca^2+^) imaging and fMRI to investigate the neural origins of fMRI-based QPPs. We uncover a robust time-locked correlation between QPPs detected by cortex-wide fluorescent WF-Ca^2+^ imaging of neural activity and QPPs of BOLD-fMRI. These data validate the hypothesis that BOLD-fMRI QPPs derive from preceding slow waves of neural activity with regional and temporal precision.

## 1. INTRODUCTION

Spontaneous brain activity measured with functional magnetic resonance imaging (fMRI) is sensitive to patterned systems-wide infraslow [0.008-0.1Hz] fluctuations, with conserved spatiotemporal characteristics across species^1–5^. As fMRI is a widely used tool for studying large-scale brain organization in both basic and clinical neuroscience, a deeper understanding of these signals has far-reaching implications for studying brain function in health and disease^6,7^. Notably, this dynamic, as opposed to static (averaged over time) treatment of fMRI data, goes beyond measures of connectivity or task/stimulus-induced responses to exploit another dimension of these rich and versatile data.

Among these dynamics, quasi-periodic patterns (QPPs) represent dominant recurring spatiotemporal motifs in fMRI data^8–12^. QPPs are typified by coordinated waves of activity between established cortical systems (e.g., default-mode and task-positive networks)^11,13–15^. These patterns, first characterized in fMRI data from rats via a robust analytical approach developed by Majeed et al. 2011, are reproducible across species^8,9,12,16^ and experimental conditions^17–19^, and have been linked to arousal^15^, neurodegenerative disease^19–21^, and neuropsychiatric conditions^22^. More broadly, these recurring patterns may reflect a repertoire of intrinsic oscillatory dynamics that organize large-scale functional connectivity and can even be used to predict activity across regions or layers of the cortex^23,24^.

However, the blood-oxygen-level dependent (BOLD) fMRI signal is an indirect measure of neuronal activity that is dependent on neurovascular coupling and prone to contamination with physiological noise (e.g., cardiovascular or respiratory)^25^. It remains possible that BOLD QPPs are generated from sequences of physiologic variation independent from neural activity. Thus, validating measures derived from BOLD-fMRI signals is critical, as is work which aims to better understand the underlying neurophysiological sources of contrast. These are well-recognized areas of focus in fMRI neuroimaging research^26–28^.

To this end, simultaneous multimodal imaging approaches, which integrate complementary sources of functional contrast, are critical despite being particularly challenging to implement (due to the high main magnetic field and limited working space within the MRI scanner)^26,28–30^. To validate fMRI-derived QPPs, simultaneous electrophysiological measures of local field potentials (LFPs) have been collected in rats and used to infer that fMRI-derived QPPs have a neural rather than an alternate source – at least relative to the point-measurements afforded by single electrodes^14,31^.

Here, we use a unique implementation of simultaneous wide-field calcium (WF-Ca^2+^) and fMRI in mice that was recently described by our group^29^. Complementing fMRI, WF-Ca^2+^ imaging is an optical technique which yields fluorescence-based measures of brain activity with high specificity and spatiotemporal resolution from a large field-of-view (FOV): the full cortical surface of the mouse^29,32–36^. Using a fluorescence-based experimental framework, the WF-Ca^2+^ imaging data in this study report on concerted pan-neuronal activity^29,32,37^. Using QPPs as a readout of large-scale brain dynamics, we apply this framework to simultaneously acquired WF-Ca^2+^ and fMRI data in healthy adult wild-type mice. The overarching aim of this work is to determine if there is a relationship between WF-Ca^2+^ and fMRI derived QPPs and by extension a solid neuronal basis for BOLD-fMRI signal derived measures of large-scale brain dynamics.

QPPs were independently identified in WF-Ca^2+^ and fMRI data, revealing a dominant spatiotemporal pattern across the cortex at both the group and subject level. The observed QPPs showed highly similar spatiotemporal organization across modalities, including coordinated activity between canonical cortical systems, in line with both human and animal fMRI literature^11,13–15^. Time courses, occurrence rates, and inter-modal as well as inter-subject (dis)similarities were evaluated. Notably, QPP occurrences aligned across modalities when accounting for an expected hemodynamic delay of ∼3–6 seconds, which we directly validated in the multimodal data^31,38–40^. Finally, we found that WF-Ca^2+^-derived patterns could successfully predict fMRI-QPPs, when accounting for the hemodynamic delay. In sum, this work establishes simultaneous WF-Ca^2+^ and fMRI as a powerful platform for quantifying brain dynamics using complementary sources of contrast. More broadly, the results herein provide clear evidence that BOLD-fMRI-derived QPPs are closely linked to neuronal activity.

## 2. RESULTS

### 2.1 Overview of multimodal data acquisition and preprocessing

Simultaneous WF-Ca^2+^ and fMRI data were obtained from anesthetized (0.5-0.75% isoflurane) mice (N = 8, 4/4 male/female) (**Methods**). To acquire WF-Ca^2+^ imaging data, mice underwent two minimally invasive surgical procedures as described previously^32^ (**Methods**). Briefly, a Ca^2+^-sensitive fluorophore (pan neuronal, hSyn-GCaMP6s^37^) was introduced at post-natal day 0 (P_0_) via transverse sinus injection (**Fig. 1A**). This procedure yields uniform cortex-wide fluorophore expression which is necessary for WF-Ca^2+^ imaging^41^ At three months of age, all mice underwent a second procedure where the skull was exposed and thinned to translucency (but left intact). A custom glass head-plate was affixed to the bone with transparent dental cement for permanent optical access to the cortical surface and immobilization during imaging^32^ (**Fig. 1B &C**). At six months of age, thirty minutes of functional multimodal data were acquired from all mice (in three 10-minute runs) using our custom multimodal imaging platform^29^ (**Fig. 1D**). Data were preprocessed and quality assessed (more below) using both the Rodent Automated BOLD Improvement of EPI Sequences (RABIES)^42^ and BioImage Suite (BIS)^43^ software packages as we have described previously^32,34,44^ (**Methods**).

**Fig. 1.**
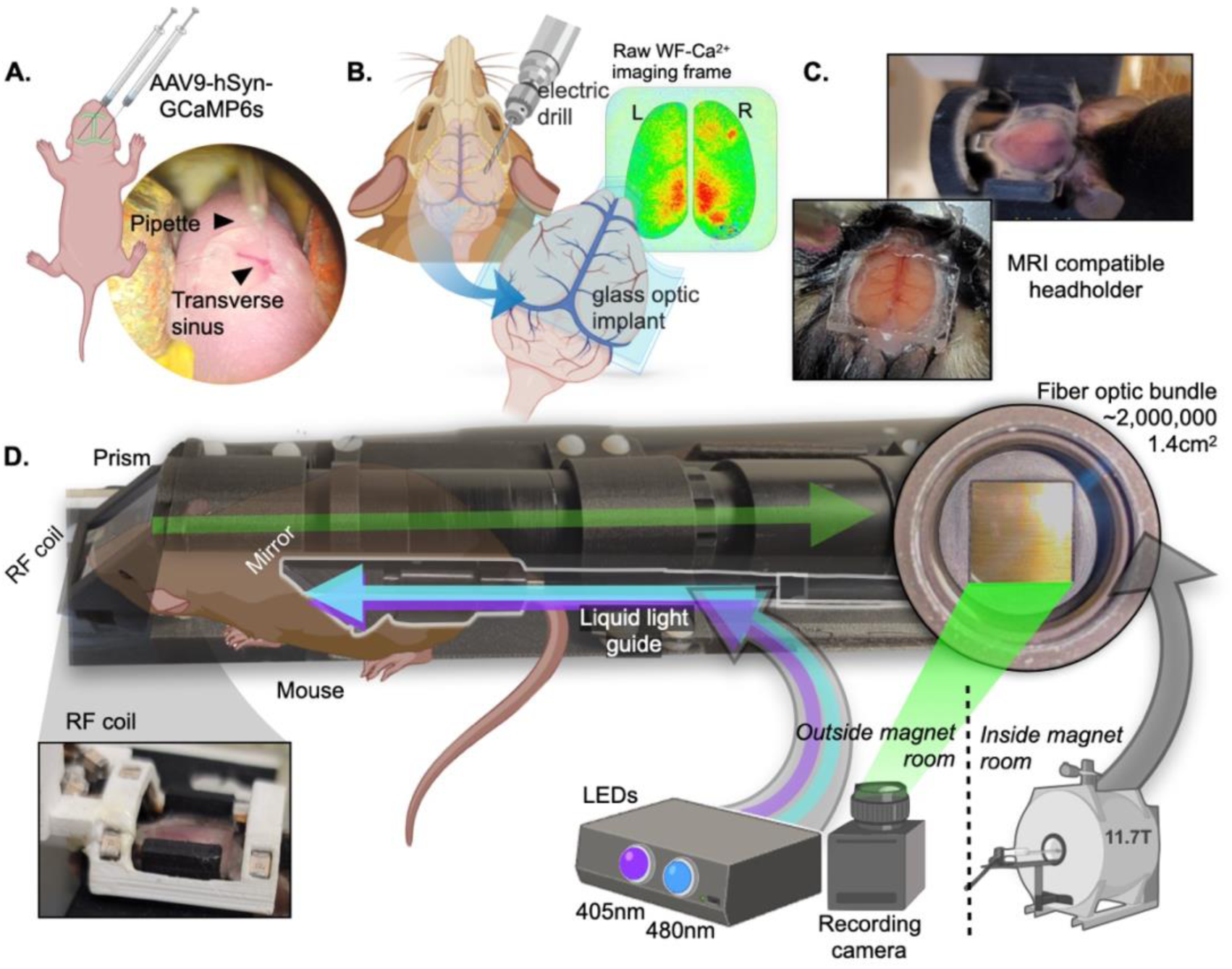
Surgical procedures and multimodal imaging platform. **A.** Transverse sinus injections performed at birth (P_0_) to introduce whole-brain fluorophore expression for WF-Ca^2+^ imaging. **B.** A schematic of the skull thinning procedure and glass head-plate attachment for optical access and head immobilization. **C.** Photographs of the glass head-plate and MRI-compatible holder. **D.** Schematic of our simultaneous WF-Ca^2+^ and fMRI apparatus. Briefly, these multimodal data were acquired synchronously using a custom-built MRI-compatible metal-free optical apparatus that is inserted into the 11cm bore of an 11.7T preclinical MRI magnet. A 2,000,000-unit fiber optic bundle relays images of the mouse cortex from within the scanner bore to the neighboring room where they are collected using two CMOS cameras (at a safe distance from the high magnetic field). A liquid light guide (15ft) is used to relay excitation light (480nm), and background illumination (405nm), for baseline correction, from the light source (also housed in the neighboring room) to magnet bore.

Data were registered to an in-house template which we have aligned with the Allen Institute Common Coordinate Framework Reference Atlas (CCfv3) and shared previously through BIS^34,45^. To compare brain activity patterns across modalities, we considered cortical regions-of-interest (ROIs) (n = 50) that are visible with both WF-Ca^2+^ and fMRI (**Fig. S1A&B**).

Time series data, for QPP^10,16^ analyses (**2.2**), were the averaged fluorescent WF-Ca^2+^ or BOLD-fMRI signals within each cortical ROI. Frequency filtering (0.008-0.2Hz)^33–35^ and global signal regression were applied to the data from both modalities (**Fig. S1C**). The WF-Ca^2+^ imaging data were temporally down-sampled to match the fMRI data (more in **Methods**). Average functional connectomes (**Fig. S1D**) as well as ROI x timepoints time series were computed for each run for both modalities. For both modalities, static average connectomes (z-scores) recapitulated stereotypical patterns including high functional connectivity between bilateral brain regions (**Fig. S1D**).

### 2.2 Group-level QPP discovery using WF-Ca^2+^ or fMRI data

A schematic of the QPP detection pipeline, as applied in this study, is illustrated in **Fig. 2** (inspired by previous illustrations^10^). For QPP detection, the within ROI averaged time series data from all mice, and runs, are used from each modality independently (**Fig. 2A&E**), i.e., QPP discovery uses either WF-Ca^2+^ or fMRI data. The algorithm is initialized using a spatiotemporal template (**Fig. 2B&F**), defined as a contiguous segment of the timeseries extracted from one mouse/run. The segment has a fixed duration which corresponds to the expected length of the QPP. Based on previous work in mice^10,16^, the expected QPP duration for our dataset is 10-seconds. This value, and the temporal resolution of the data, are used to compute an expected ‘phase length’ (PL) [1] (reproduced from^10^).

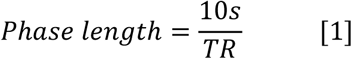

**Fig. 2.**
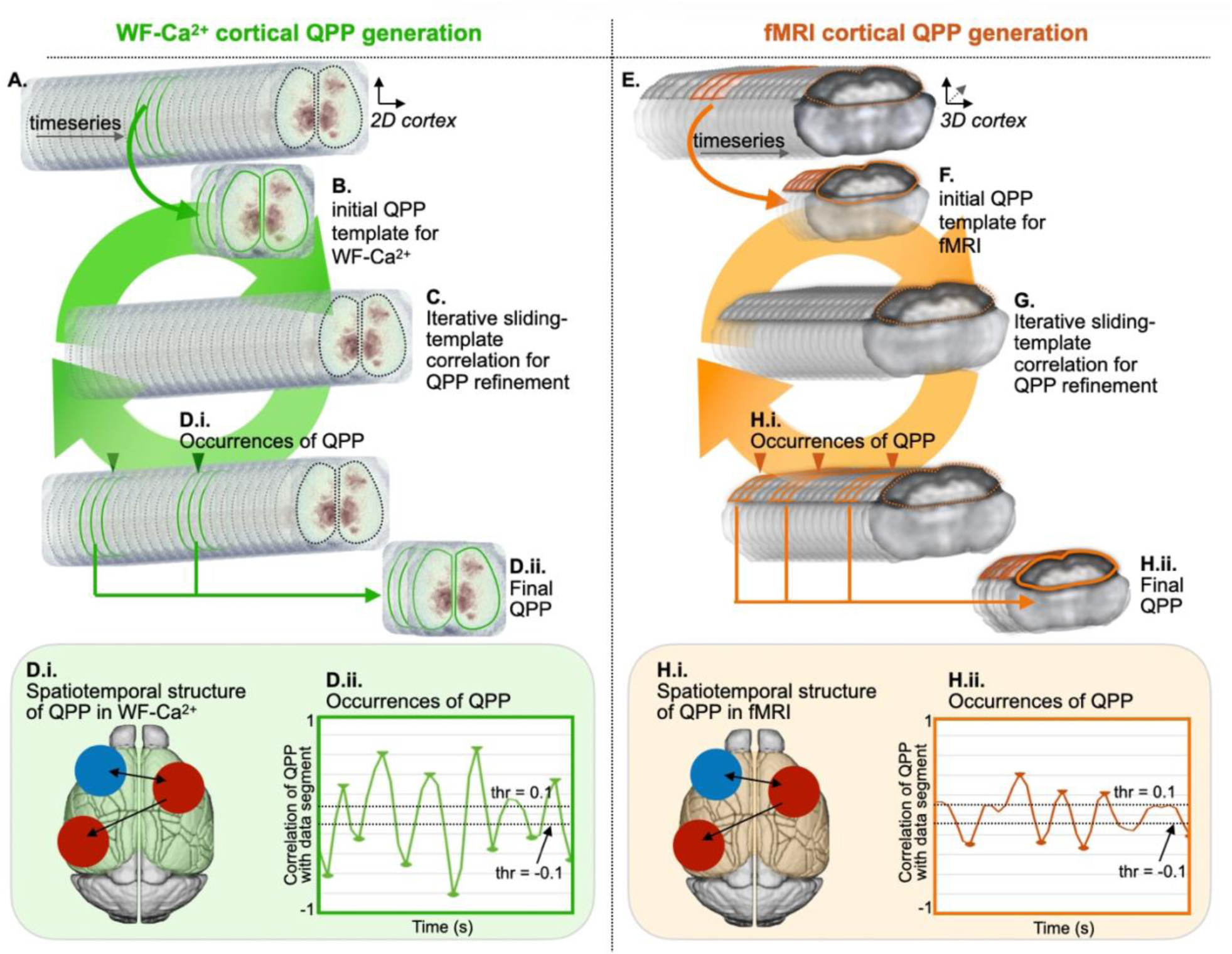
QPP discovery and quantification using WF-Ca^2+^ or fMRI data. Schematic of the QPP detection pipeline applied to the simultaneously acquired WF-Ca^2+^ (left, green) or fMRI (right, orange) data. For each imaging modality, timeseries data **(A&E)** from 50 *a priori* cortical ROIs (**Fig. S1**) are used. An example randomly selected initialization template **(B&F)** is iteratively refined by sliding-window correlation across the timeseries (all mice/runs) **(C&G)** to identify QPP occurrences across each run (**D.i&H.i**; example occurrences highlighted with triangles). A final QPP (**D.ii&H.ii**) is identified when iteratively refined templates are compared to identify the one with the highest summed correlation across occurrences. Occurrences of the QPP are identified in the dataset by correlating the QPP with the timeseries and applying a cut-off: r ≧ 0.1 for in-phase, and r ≦ −0.1 for anti-phase **(D.ii&H.ii)**.

Here, QPP duration (10-seconds), and repetition time (TR) 1.8-seconds (**Methods**), corresponds to a PL of 5.5 frames. Because this number must be an integer, we use a PL of 6. PL of 6 defines half of the QPP cycle (in-phase); the second half of the QPP cycle is also considered (anti-phase) for a total of QPP length of 12 frames (timepoints). Using a PL of 5 yields near-identical results (data not shown). With this, QPP discovery follows a 4-step framework:

Step 1. An initialization template (frames 1-6 for in-phase) is correlated with all valid segments of the timeseries of each mouse/run yielding an iterative sliding-template Pearson correlation trace (**Fig. 2C&G**).

Step 2. Candidate occurrences are identified as local maxima in the correlation trace when correlation with the initialization template exceeds a threshold (|0.1| for iterations 1, 2, & 3, and |0.2| thereafter, refer to Step 3). Overlapping detections are avoided by imposing the criterion that occurrences must be separated by at least one template length (**Fig. 2D.i&H.i**).

Step 3. A refined template is derived by averaging across candidate occurrences (across mice and runs, for both in-phase and anti-phase) (**Fig. 2D.ii,H.ii**). The refined template is then reapplied to the dataset and Steps 1-3 repeated iteratively (to a maximum of 20) or until convergence (QPP does not change).

Step 4. This process is then repeated beginning with a new initialization template (2-7, 3-8, etc.) selected initialization template. The initialization template was selected by sliding a window of 12 frames across the whole time course, which is an exhausted search that covers all candidate templates. The algorithm terminates when all possible initialization templates have been tested (robust mode^10^). The resulting (fully refined and stable) templates are then compared and the solution, or final QPP, is identified as the refined and stable template with the greatest summed correlation across occurrences within the dataset.

As a control, either ROI labels or imaging frames were shuffled randomly within each run; in both cases, the spatio-temporal structure of the pattern was not recovered (data not shown).

The spatiotemporal QPP discovered, i.e., regions that are either co-activated, or deactivated, is defined by the average of all individual occurrences **(Fig. 2D.i&H.i**). Occurrences of the QPP can also be examined by extracting the timepoints at which the QPP appears using the same definition for occurrence as used for QPP discovery (here, |0.1|, see Step 2) (**Fig. 2D.ii&H.ii**).

### 2.3 Spatiotemporal characteristics of the QPP are preserved across modalities

The spatiotemporal characteristics of the group-derived QPP (across all mice/runs), discovered using either fluorescent WF-Ca^2+^ imaging or BOLD-fMRI signals, are shown in **Fig. 3** (left, right, respectively). Specifically, **Fig. 3A** depicts the spatial distribution of cortical involvement at successive timepoints across the QPP; these results are also shown as a 3-D rendering on the cortical surface in **Fig. 3D** (and as a time-lapse sequence in **QPP_media.mov**). Using data from either imaging modality, the QPPs spanned 12-timepoints (each corresponding to TR = 1.8 seconds, **Methods**) and are composed of two distinct halves (6 in-phase, and 6 anti-phase). As depicted in **Fig. 3A&D**, the in-phase half is characterized by motor (MO) and somatosensory (SS) areas being ‘up’ (positive, red, during in-phase) and auditory (AUD), visual (VIS), and retrosplenial (RSP) areas being ‘down’ (or negative during in-phase) followed by a mid-point transition (timepoint 6) during which there is a reversal as the anti-phase half begins: MO and SS areas transition ‘down’ (negative) while AUD, VIS, and RSP areas transition ‘up’ (positive).

**Fig. 3.**
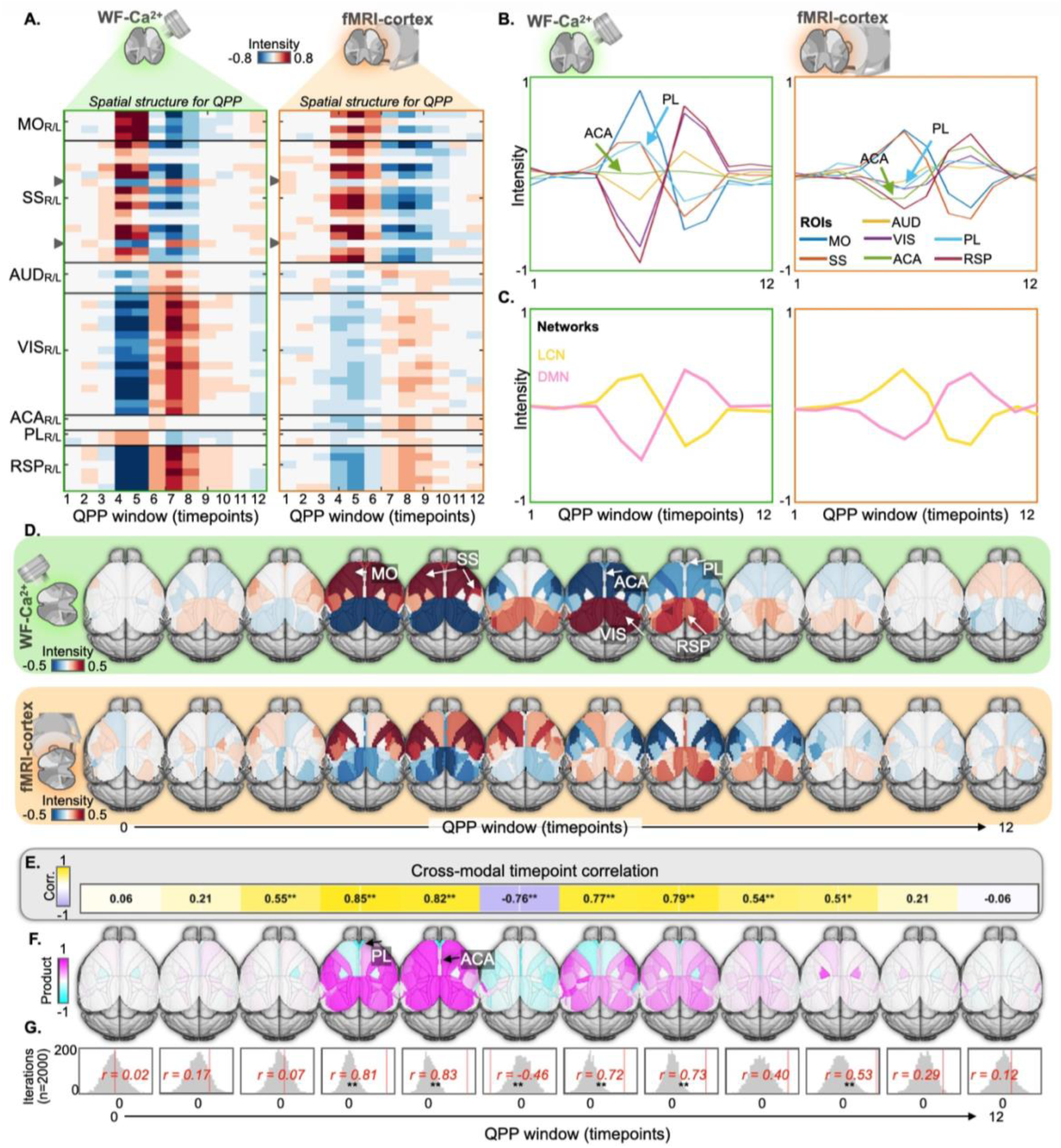
Spatiotemporal properties of group-derived QPPs and cross-modal correspondence. **A.** Spatiotemporal tructure of the QPPs discovered using WF-Ca^2+^ (green, left) or fMRI (orange, right) data and 50 *a priori* cortical ROIs **Fig. S1A**. SS-trunk is indicated by grey triangles as an outlier amongst all other SS regions. **B.** Profile of the QPPs across ROIs for WF-Ca^2+^ (left), and fMRI (right). **C.** Profiles of the QPPs averaged across ROIs in two representative functional networks: DMN (pink) and LCN (yellow). **D.** Surface rendering of QPPs at successive timepoints (WF-Ca^2+^ along the top row, and fMRI along the bottom row). **E.** Cross-modal timepoint-resolved correlation (Pearson’s, Bonferroni corrected) between QPPs discovered with WF-Ca^2+^ and fMRI data. **F.** Surface rendering of z-scored WF-Ca^2+^ × fMRI QPPs at each timepoint to highlight cross-modal QPP coherence across ROIs and time. **G.** Cross-modal timepoint-resolved correlation Pearson’s) between QPPs discovered with WF-Ca^2+^ and fMRI data *after* z-scoring (i.e., as in **E.** with the addition of z-coring) alongside a null distribution generated by shuffling ROIs in the QPP generated from the WF-Ca^2+^ imaging data 2,000 iterations). Observed correlations indicated in red. MO: motor cortex, SS: somatosensory cortex, AUD: auditory cortex, VIS: visual cortex, ACA: anterior cingulate area, PL: prelimbic area, RSP: retrosplenial cortex, DMN: default-mode network, LCN: latero-cortical network, ROIs: regions-of-interest, R/L: right/left. *p <0.05, **p <0.01, ***p <0.001.

Examples of the WF-Ca^2+^ and fMRI derived QPPs within cortical ROIs are shown in **Fig. 3B**, following the Allen Atlas CCfv3^45^ as summarized in **Fig. S1**. Note that the data in **Fig. 3** show modality-specific time courses of the QPPs. Thus, timepoint correspondence across modalities is in terms of the QPP ‘structure’ – not *when* the QPP occurs in one modality relative to the other. Relative timing is considered below.

In-line with previous work^11,46^, which has largely examined fMRI data, a hallmark of QPPs is captured by imposing canonical functional network definitions (**Fig. S1A**). Specifically, cortical ROIs within the default-mode network (DMN): RSP, anterior cingulate cortex (ACA), and prelimbic (PL) cortex, when grouped together, exhibit a common pattern (‘down’ for in-phase followed by ‘up’ for anti-phase), which is anticorrelated with the pattern shown by the lateral cortical network (LCN or task-positive network), comprised of: SS and MO regions (**Fig. 3C**).

While this network-level pattern was well recapitulated in the fMRI data (**Fig. 3C**, right) – and observed in the averaged WF-Ca^2+^ imaging data (**Fig. 3C**, left) – there were some brain regions, in the later modality, that did not follow this expected pattern. Specifically, the ACA (**Fig. 3B**, left, green line), part of the DMN, showed no pattern, while the PL cortex **(Fig. 3B**, left, cyan) – also part of the DMN – showed the opposite to expected pattern. Further, in-line with previous work^25^, an anticorrelation between SS and VIS areas was observed (**Fig. 3B**).

Another observation is that the SS-trunk ROI (notably, both left and right), in the QPPs recovered using either imaging modality, showed the opposite to expected pattern, i.e., ‘down’ for in-phase and ‘up’ for anti-phase, compared to all other SS ROIs (**Fig. 3A**, SS-trunk right/left indicated by grey triangles). Nevertheless, despite some inter-modal differences and ROI-specific findings, the overall spatiotemporal characteristics of the QPPs recovered using either WF-Ca^2+^ or fMRI data were remarkably consistent across modalities and showed established characteristics of mouse cortical functional organization^11,16^. This included notable bilateral symmetry in the QPPs, illustrated more clearly in **Fig. S2**, where the ROIs are separated by hemisphere.

### 2.4 Mismatch across modalities at the mid-point transition between in-phase and anti-phase

To quantify agreement between group-derived QPPs derived from WF-Ca^2+^ and fMRI data, a vector of ROI × intensity within each of the 12 timepoints within the QPPs were correlated across modalities (**Fig. 3E**). This revealed strong cross-modal agreement at timepoints 4-5 and 7-8 (r = 0.85/0.82 for timepoints 4/5; and r = 0.77/0.79 for timepoints 7/8, p < 0.01, corrected). In contrast, timepoint 6 exhibited a marked anticorrelation between the QPPs derived from WF-Ca^2+^ versus fMRI data (r = −0.76, p < 0.01, corrected), suggesting a mismatch in the mid-point transition between the in-phase to anti-phase halves of the QPPs across modalities. This mismatch is also evident, upon closer inspection, in the 3D-renderings in **Fig. 3D**, and in the ROI × timepoint matrix in **Fig. 3A**.

To quantify these dynamics at the level of ROIs, the QPPs detected with WF-Ca^2+^ and fMRI data were compared by taking a cross-modal product (WF-Ca^2+^ × fMRI) at each timepoint after z-scoring (**Fig. 3F**). Significant correspondence between QPPs across modalities was again very apparent during timepoints 4-5 as well as 7-8 with a mismatch during the transition/mid-point at timepoint 6. Comparisons to data where the ROIs were shuffled (after z-scoring) at each timepoint within each modality (**Fig. 3G**) confirmed significant positive correlations for timepoints 4-5, 7-8, and 10 (e.g., r = 0.81 for timepoint 4, p < 0.01, corrected), while timepoint 6 showed a significant anticorrelation (r = −0.46, p < 0.01, corrected).

Overall, these findings illustrate the expected more rapid dynamics in the fluorescent WF-Ca^2+^ imaging signal relative to the BOLD-fMRI signal.

### 2.5 Occurrences of WF-Ca^2+^ or fMRI derived QPPs in source-matched data

Occurrences of the WF-Ca^2+^ and fMRI derived QPPs are shown in **Fig. 4**. For each subject, group-derived QPP occurrences are identified by sliding the corresponding modality-specific QPP across the time course and correlating with each 12 timepoint segment. Correlation strengths, between the occurrence time courses (WF-Ca^2+^ imaging in green or fMRI in orange) and each corresponding modality-specific QPP, are shown in **Fig. 4A** where positive values correspond to in-phase occurrences (r > 0.1, triangles) and negative values correspond to anti-phase occurrences (r < −0.1, circles). As in QPP discovery (**2.2**, step 2), overlapping detections are avoided by imposing the criterion that occurrences must be separated by at least 12 timepoints.

**Fig. 4.**
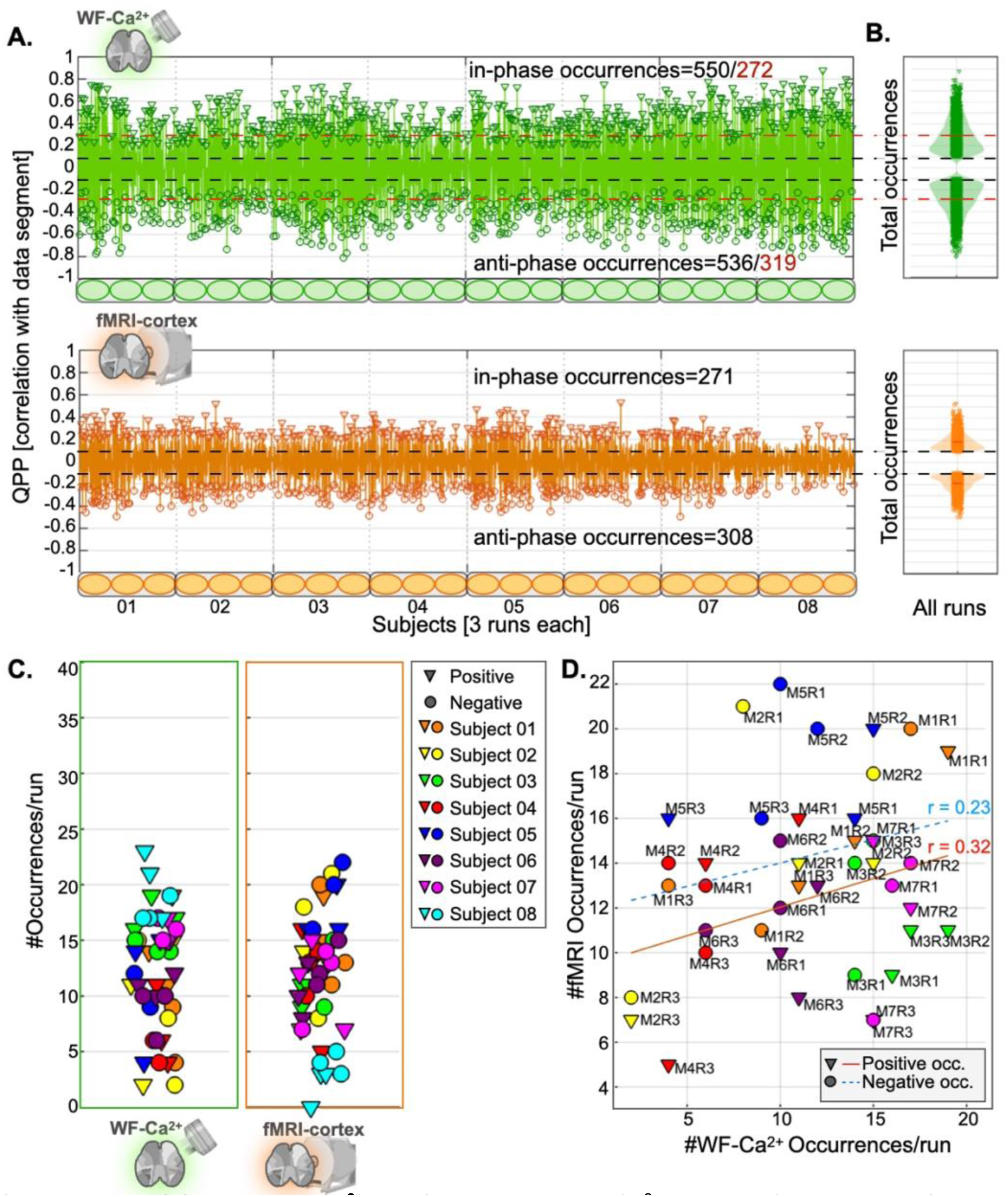
Occurrences of QPPs in WF-Ca^2+^ and fMRI data. **A.** WF-Ca^2+^ (top) and fMRI (bottom) QPP occurrences (triangles/circles denote in-/anti-phase). Subjects (grey boxes) and runs (green/orange ovals, for WF-Ca^2+^ and fMRI, respectively) are concatenated. Correlation of modality-specific QPPs with all data segments within the source data; positive values correspond to in-phase QPP occurrences and negative values correspond to anti-phase occurrences. Dashed black lines indicate the detection threshold |0.1|. Dashed red lines, for WF-Ca^2+^ imaging data, indicate the detection threshold |0.3|. **B.** Violin plots: kernel density distribution of values exceeding |0.1| for WF-Ca^2+^ imaging (top) and fMRI (bottom) data. Individual data points are overlaid as a scatterplot. **C.** Total number of occurrences above the mismatched thresholds (|0.3| and |0.1|, for WF-Ca^2+^ and fMRI, respectively) per run for each mouse. In-phase occurrences are denoted with triangles. Anti-phase occurrences are denoted with circles for WF-Ca^2+^ (left) and fMRI (right) data. **D.** Cross-modal correlation of occurrences for in-phase and anti-phase occurrences (using the mismatched thresholds). Regression lines for in-phase (red solid) and anti-phase (dashed cyan) occurrences are shown despite not being statistically significant (p-values = 0.16 and 0.31, respectively).

Across subjects, and runs, both modalities exhibited frequent WF-Ca^2+^ and fMRI derived QPP occurrences in the respective source data, **Fig. 4A&B**. In-phase and anti-phase occurrences included a range of correlation strengths between the WF-Ca^2+^ or fMRI derived QPPs and the source data, **Fig. 4B**. Overall, the WF-Ca^2+^ imaging derived QPP showed higher correlation strengths with the WF-Ca^2+^ imaging data segments and more frequent occurrences (in-phase and anti-phase) than the fMRI derived QPP and fMRI data. When the occurrence threshold applied to the WF-Ca^2+^ imaging data (and corresponding QPP) was increased, to |0.3|, the number of occurrences was nearly matched across modalities (**Fig. 4A**, red dashed lines and red text).

Although not statistically significant, the number of occurrences per run showed a positive correlation across modalities which improved when mismatched detection thresholds were applied (**Fig. 4D**). For this comparison subject 08 was removed (more below), r = 0.14/0.25 (p = 0.53/0.28) for in-phase/anti-phase, respectively, using a |0.1| threshold for both modalities; r = 0.32/0.23 (p = 0.16/0.31 for in-phase/anti-phase respectively) using |0.3| and |0.1| for WF-Ca^2+^ and fMRI, respectively. This indicates that although it may be easier to recover QPPs from WF-Ca^2+^ imaging data, there is a relationship across modes such that QPPs, derived from either modality, occur with related frequencies.

Subject 08 was identified as a clear outlier when occurrence rates were compared across imaging modalities. This subject showed the highest number of WF-Ca^2+^ imaging derived QPP occurrences alongside the lowest number of fMRI derived QPP occurrences (in-phase and anti-phase, **Fig. 4C**). This subject also showed high correlation strengths between the WF-Ca^2+^ imaging derived QPP and corresponding data whilst the fMRI derived QPP showed low correlation strengths. To assess whether these observations could be attributed to data quality, we examined the temporal signal-to-noise ratio (tSNR), **Fig. S3**, head motion, **Fig. S4**, and specific connectivity, **Fig. S5**, of subject 08 relative to the other 7 mice, see **Methods**. No differences in tSNR (**Fig. S3**), motion (**Fig. S4**), or WF-Ca^2+^ specific connectivity (**Fig. S5A**) across subjects accounted for these observations. The only measure which set subject 08 apart was low specific connectivity in the fMRI data (**Fig. S5B**).

### 2.6 Subject specific QPPs from WF-Ca^2+^ and fMRI data

In high-quality data, QPPs can be derived from small amounts of data collected from individual subjects^10^. To assess this possibility in our multimodal data, we repeated the QPP discovery pipeline (**2.2**, **Fig. 2**) using either WF-Ca^2+^ or fMRI data from each mouse independently. WF-Ca^2+^ or fMRI single-subject derived QPPs from each mouse were compared to the modality-matched group-derived QPPs, as well as across subjects and modalities. These results are summarized in matrices, **Fig. 5**, where the QPPs (as in **Fig. 3A**) were ‘unwrapped’ into a vector and correlated: cross-subject, subject-to-group, and cross-modality. Of note, the quantifications within this section are intended as exploratory assessments of inter-subject variability and do not impact the main results of this work, for which all subjects are retained.

**Fig. 5.**
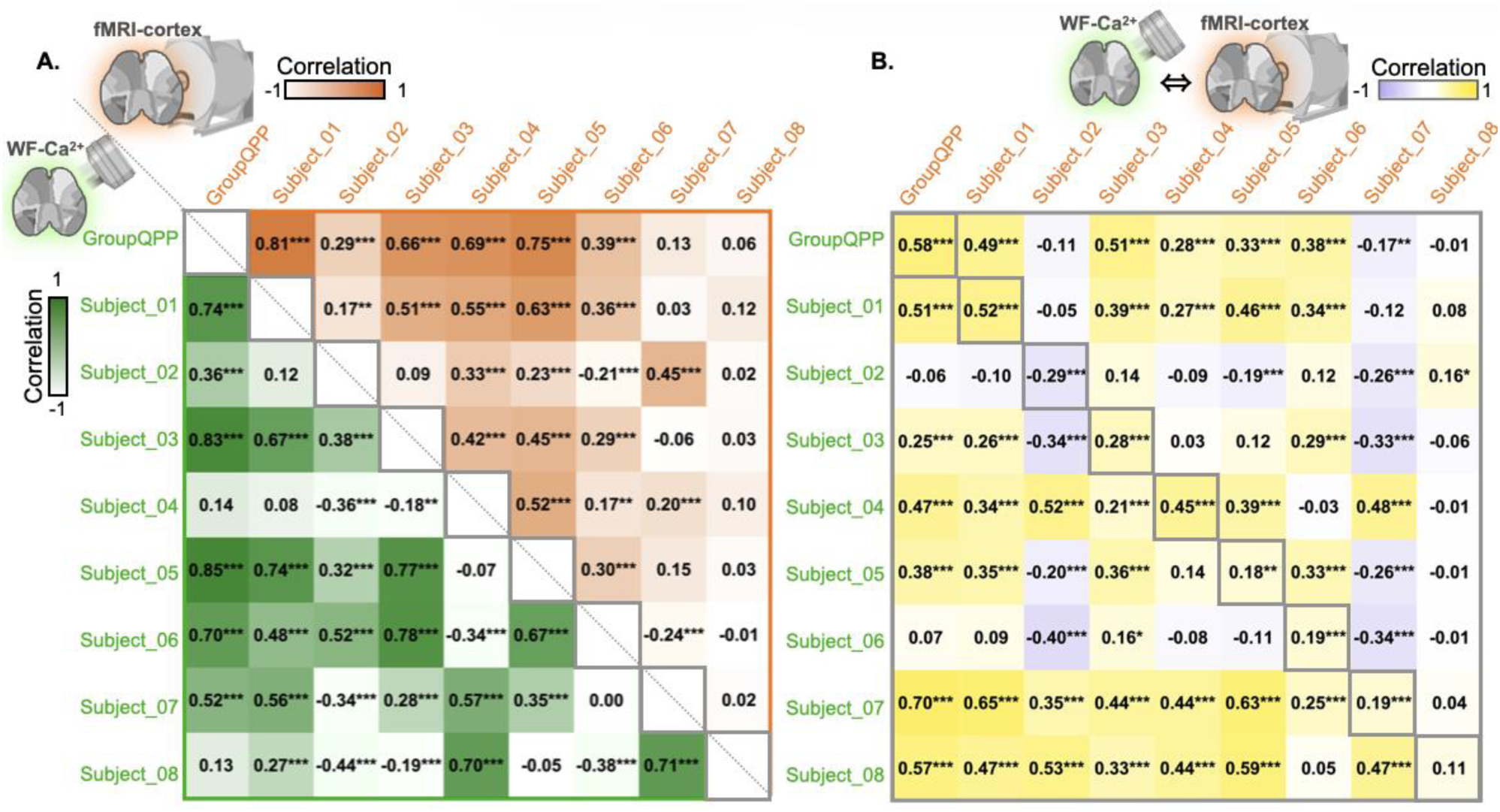
Within and cross-modality group and subject-derived QPP comparison. Matrices compare group-(rows/columns 1) and subject- (rows/columns 2-9) derived QPPs within- (**A**) and across (**B**) modalities. **A.** WF-Ca^2+^ (green hues), and fMRI (orange hues) derived QPPs are compared within modality. **B.** WF-Ca^2+^ and fMRI derived QPPs are compared across modalities within (diagonal) and across (off-diagonal) subjects. Correlation coefficients (Pearson’s r, Bonferroni corrected) are overlaid on each cell. Asterisks denote statistical significance after correction, *p < 0.05, **p < 0.01, ***p < 0.001).

Most subject-specific QPPs, derived from WF-Ca^2+^ or fMRI data, showed good agreement across mice within modality: r = 0.43 ± 0.31, excluding subjects 04 & 08, and r = 0.33 ± 0.18, excluding subjects 07 & 08, for WF-Ca^2+^ and fMRI derived QPPs respectively, mean ± standard deviation (STD), (**Fig. 5A**). Subjects 04 & 08 as well as 07 & 08, for WF-Ca^2+^ and fMRI data respectively, were excluded in this specific-analysis based on low correlations between their subject-specific QPPs and the group-derived QPP (row/column 1). Importantly, both subjects 04 and 07 showed ‘specific’ functional connectivity (within the bottom-right quadrant, **Fig. S5**), for fMRI, and no removed frames based on motion (data not shown). When subject-specific QPPs from subjects 04 & 08 were examined (**Fig. S6**), the WF-Ca^2+^ derived QPPs showed a shift in their temporal pattern where the QPP appeared to have a delayed beginning and end (black vs grey triangles in **Fig. S6**), as well as later inflection point or switch between halves (at timepoint 7 rather than 6, see **2.3**). In-line with this observation, subjects 04 & 08 also showed a high correlation between their subject-specific QPPs derived from the WF-Ca^2+^ imaging data (r = 0.70, p < 0.001, corrected). On the other hand, the subject-specific QPPs derived from the fMRI data for subjects 07 & 08 did not show a clear spatiotemporal pattern.

Across modalities, the majority of subjects showed good within-subject agreement between subject-specific WF-Ca^2+^ and fMRI derived QPPs (r = 0.27 ± 0.15, excluding subject 02, **Fig. 5B**, matrix diagonal). The group-derived QPPs, from WF-Ca^2+^ and fMRI data, were also correlated (r = 0.58 p < 0.001, corrected). Likewise, across subjects and modalities, subject-derived QPPs were correlated: r = 0.37 ± 0.18, excluding subjects 02 & 06, bottom half columns 2-9, and r = 0.26 ± 0.16, excluding subjects 02, 07 & 08, top half rows 2-9. As above, subjects 02 & 06 for fMRI, as well as 02, 07 & 08, for WF-Ca^2+^ were excluded in this specific analysis based on low correlations between the subject-specific QPPs and the group-derived QPP (row/column 1). Importantly, subject 06 showed functional connectivity specificity well within the ranges of ‘high specificity’ across both modalities (**Fig. S5**), as well as no frames discarded based on motion (data not shown).

Altogether, these results show that: (1) Subject-specific QPPs can be derived from most subjects using either WF-Ca^2+^ or fMRI data which are in good agreement with the group-derived QPP. (2) When subject-specific QPPs derived from WF-Ca^2+^ imaging data deviate from the group-derived QPP, it may be an indication that the temporal characteristics of the QPP in that subject are skewed. Whereas when the subject-specific QPP derived from fMRI data deviate from the group-derived QPP, it is more likely due to a lack of a recoverable pattern in the data. (3) Most subject-specific QPPs show good agreement across modalities both within and across subjects. As expected, when subject-specific QPPs from one modality are not correlated with the group-derived QPP from the second modality, the cross-modal correlation of that subjects’ QPP(s) with itself and others is low.

### 2.7 Cross-modal temporal alignment of WF-Ca^2+^ and fMRI derived QPPs

Neurovascular coupling operates with a well-established hemodynamic delay^47^. Evidence of this delay in our multimodal data was noted above in the mismatch, at timepoint 6 (**Fig. 3**), between WF-Ca^2+^ and fMRI at the transition point between in-phase and anti-phase halves of the QPPs derived from each modality. Here, we quantify the temporal shift between fluorescent WF-Ca^2+^ and BOLD-fMRI signals using cross-modal correlation with systematically added temporal lags to identify the ‘best match’ between modalities (**Fig. 6**). For this analysis we use the occurrence time courses from each mouse/run which are derived by correlating the modality-specific group QPP with the source-signal time course in a maximally overlapping framework, see above (**2.4**) and **Methods**.

**Fig. 6.**
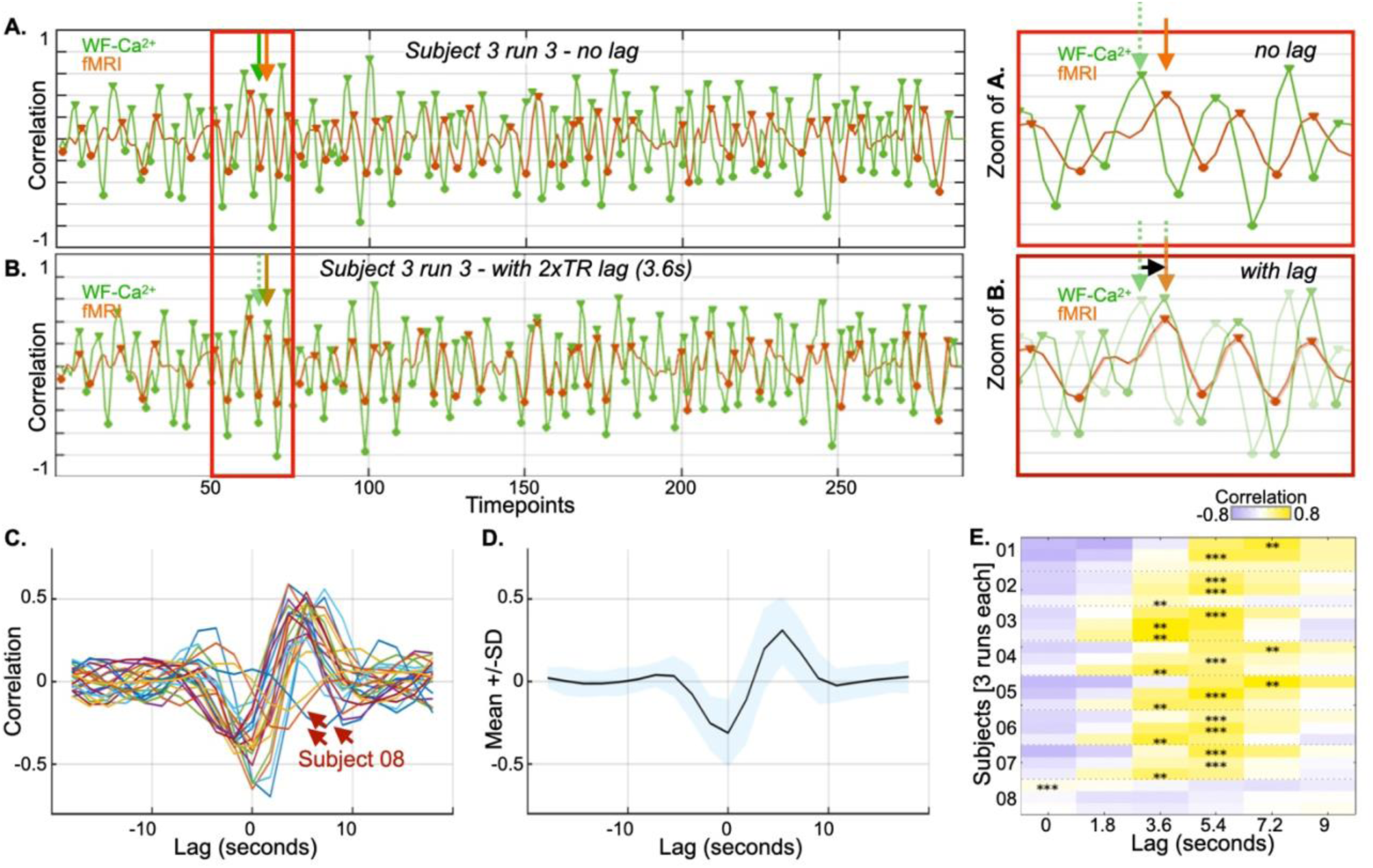
Cross-modal temporal alignment of QPP between fMRI and WF-Ca^2+^ data. **A.** Representative QPP template-correlation time courses from a single run (Subject 3, run 3) shown without temporal lag correction. Orange traces indicate fMRI-cortex and green traces indicate WF-Ca^2+^ cortex. The y-axis represents correlation with the QPP template; filled markers denote detected phase and antiphase events. The boxed region (right) highlights misalignment between modalities prior to lag correction. **B.** Same run after applying a temporal lag to the BOLD signal. Alignment of QPP occurrences across modalities improves substantially, as illustrated in the zoomed panel (right), where corresponding peaks and troughs show enhanced temporal overlap. **C.** Cross-modal correlation as a function of temporal lag for all mice and runs, demonstrating a consistent peak at positive lags. **D.** Group-averaged cross-modal correlation curve (mean ± SD), revealing maximal correspondence at ∼3.6-5.4 s. **E.** Heatmap of cross-modal QPP correlation values across runs for discrete lag values (0, 1.8, 3.6, and 5.4 s). Best lags (highest correlation) are indicated by significant p values, with *p<0.05, **p<0.01 and ***p<0.001.

Representative WF-Ca^2+^ and fMRI (green and orange, respectively) occurrence time courses (subject 03, run 3), are shown before and after applying a temporal lag of 3.6-seconds or 2TRs to the WF-Ca^2+^ imaging data (**Fig. 6A&B**). Without this lag, QPP occurrences are clearly misaligned across modalities with a total of 20 overlapping ‘peaks’ (14 in-phase and 6 anti-phase) and a cross-modal correlation r = −0.19 (p < 0.01) (**Fig. 6A**). After applying a temporal shift (of 3.6-seconds or 2TRs), alignment between the WF-Ca^2+^ and fMRI occurrence time courses improves (**Fig. 6B**). The number of overlapping ‘peaks’ increases to 51 (28 in-phase and 23 anti-phase), and the correlation between the occurrence time courses increases r = 0.59 (p < 0.001). As a comparison, an example run from subject 08 is shown in **Fig. S7**. A non-significant correlation r = 0.01 (p = 0.8) is found with no lag (7 overlapping in-phase peaks, 11 overlapping anti-phase peaks) and a significant anti-correlation r = −0.18 (p < 0.01) is found with a 2TR lag (6 overlapping peaks for in-phase and 6 for anti-phase).

To find the optimum lag time for the group (N = 8), we computed cross-modal correlation as a function of temporal offset for all mice/runs, **Fig. 6C**. Cross-correlation curves consistently showed a maximum at a positive lag time, indicating that BOLD-fMRI follows fluorescent WF- Ca^2+^ signals, as expected. Across 24 runs, the optimal lag was most commonly observed at 3.6-7.2 seconds, with 5.4 seconds representing the modal peak. Peak cross-correlation values were moderate to high in most runs (up to r = 0.590), consistent with strong alignment between modalities, although one mouse (subject 08) showed weak correspondence across all runs (r = 0.013-0.074). The group-averaged correlation curve (**Fig. 6D**; mean ± SD) shows a maximum at a lag time of ∼5-seconds. This lag dependence was significant at the group level, as shown by repeated-measures ANOVA across the 0-9 second window (F(5,115) = 26.12, p < 0.001). Bonferroni-corrected post hoc comparisons demonstrated that correlation strength at 3.6, 5.4, and 7.2 seconds exceeded that observed at 0 seconds, with the largest effect at 5.4 seconds. Correlations at 3.6 and 5.4 seconds were not significantly different from one another, nor were those at 3.6 and 7.2, suggesting a broad peak.

Across runs (**Fig. 6E**), most inter-modal occurrence time courses showed a maximum correlation strength at a lag of 5.4-seconds (10/24 runs) while a smaller fraction (7/24 runs) showed a maximum at 3.6-seconds or 7.2 seconds (3/24). Maximum correlation strengths ranged between 0.18 and 0.59. In clear opposition to the trend evident in N = 7 mice, three runs (all from subject 08) showed a maximum (but near zero) inter-modal occurrence time course correlation strength at a zero-lag time.

In sum, these findings indicate that the strongest coupling between spontaneous patterned fluorescent WF-Ca^2+^ and BOLD-fMRI signals occurs within a 3.4-7.2 second lag, consistent with the hemodynamic delay between neural activity and the BOLD-response^7,39^.

### 2.8 Cross-modal QPP swap: applying the WF-Ca^2+^ QPP to fMRI data, and *vice versa*

To determine whether the QPP discovered using one imaging modality captures similar dynamics in the second modality, we performed a cross-modal ‘QPP-swap’ analysis^19^. In specific terms, the group QPP from the WF-Ca^2+^ imaging data (**Fig. 3A**, left) was applied to the fMRI data using the same procedure as above (**2.3** and **2.5**) to detect ‘QPP-swap’ derived occurrences^19^ (**Fig. 7A**, top). Similarly, the group QPP from the fMRI data (**Fig. 3A**, right) was applied to the WF-Ca^2+^ imaging data (**Fig. 7A**, bottom), to detect ‘QPP-swap’ derived occurrences. This procedure results in two ‘QPP-swap’ derived occurrence time courses in addition to the original occurrences (QPP in source-matched data) reported in **Fig. 4A**.

**Fig. 7.**
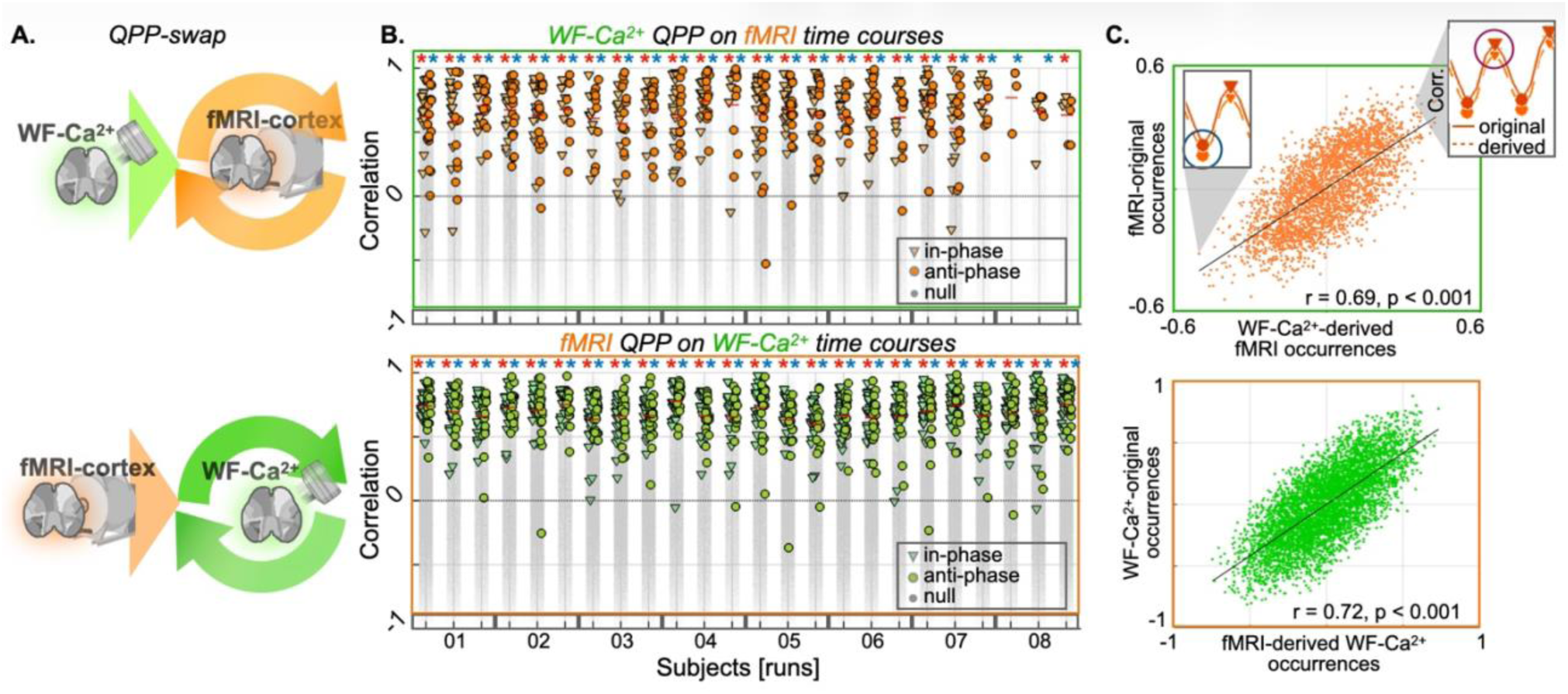
Cross-modal QPP-swap. **A.** Schematic of QPP-swap analyses using the same sliding-correlation procedure used for the original analyses (**2.3/2.5**). Top: WF-Ca^2+^ QPP applied to fMRI data to detect ‘QPP-swap’ derived fMRI occurrences. Bottom: fMRI QPP applied to WF-Ca^2+^ data to detect ‘QPP-swap’ derived WF-Ca^2+^ occurrences. **B.** Segment-wise Pearson’s correlation (Bonferroni corrected) between original and ‘QPP-swap’ derived occurrence time courses (threshold r, |0.1|). Positive occurrences are indicated with reversed-triangles, negative occurrences are indicated with circles. Gray points indicate correlations obtained from a null distribution generated by randomly permuting the temporal order of the original occurrence trace within each run (2,000 iterations). Red horizontal bars indicate the mean correlation across positive and negative occurrences within each run. Significant difference from the null (Pearson’s r, p<0.05, Bonferroni corrected) indicated for both positive and negative occurrences in red/blue respectively, above each run. Top: green outline indicating WF-Ca^2+^-QPP, orange hues indicating ‘QPP-swap’ derived fMRI occurrences. Bottom: orange outline indicating fMRI-QPP, green hues indicating ‘QPP-swap’ derived WF-Ca^2+^ occurrences. **C.** Cross-modal relationship between original and ‘QPP-swap’ derived occurrence time courses for fMRI data (top, orange dots, green outline) and WF-Ca^2+^ data (bottom, green dots with orange outline). Insets: detail of correlation for positive and negative occurrences. *p<0.05, corrected.

Each occurrence identified in the original occurrence time course (r ≧ |0.1|, **Fig. 4A**) was used to isolate a 6-frame epoch in both the original and ‘QPP-swap’ derived occurrence time course (for both modalities). The Pearson correlation of each 6-frame epoch was computed between the original and QPP-swap derived occurrence time course (**Fig. 7B**). The 6-frame epoch length corresponds to the duration of half a cycle of the QPP, 6 frames for in-phase, and 6 for anti-phase (**Fig. 3A**). A null distribution was generated by randomly permuting the original occurrence time courses within each run (2,000 iterations). Across mice/runs original and QPP-swap derived occurrence time courses were modestly-to-highly correlated and well-above the null distribution (p < 0.05, Bonferroni corrected) for both modalities, for most runs in the case where WF-Ca^2+^-QPP was applied to fMRI data (**Fig. 7B**, top), and in all runs in the case where fMRI-QPP was applied to WF-Ca^2+^ data (**Fig. 7B**, bottom). Specifically, for the former case, 22/24 runs for in-phase occurrences and 23/24 runs for anti-phase occurrences showed significant difference from the null (p < 0.05, Bonferroni corrected). Even subjects that had very few occurrences, e.g., fMRI-subject 08, still had high correlations between original and QPP-swapped occurrence time course epochs, besides two runs of subject 08 for in-phase, and one run from subject 08 for the anti-phase which either did not present any identified occurrences, or did not survive correction (**Fig. 7B**, top).

Finally, original and ‘QPP-swap’ derived occurrence time courses were compared by generating a scatterplot of original versus derived occurrence time course correlation strengths at every timepoint (across mice/runs). In other words, the sliding Pearson correlation values (**2.2** step 1) for swapped versus original (source-data) QPPs and WF-Ca^2+^ or fMRI data (**Fig. 7C**, top and bottom respectively). The scatterplots generated by applying the group WF-Ca^2+^ QPP to the fMRI data as well as the reverse (group fMRI QPP to the WF-Ca^2+^ data) both showed a strong positive correlation: r = 0.69 and 0.72, p < 0.001 corrected, respectively (**Fig. 7C**, top/bottom respectively).

Altogether, these results demonstrate that QPPs derived from WF-Ca^2+^ or fMRI data capture temporal features that are present in the other modality.

### 2.9 WF-Ca^2+^ QPP occurrences predict the spatiotemporal structure of fMRI QPPs with a temporal delay

To test whether WF-Ca^2+^-QPP occurrences could predict the spatiotemporal structure of the fMRI QPP, we reconstructed a ‘predicted’ fMRI QPP (fMRI-QPP predicted) using WF-Ca^2+^ occurrences time series, shifted by a delay of *k* frames (*k* = 0, 1, 2…9; 1 frame = 1 TR) and compared each predicted QPP to the original group fMRI QPP. We considered positive and negative occurrences from the ‘mismatched’ correlation threshold reported above for WF-Ca^2+^(see **2.5**) where the number of occurrences across modalities was highly similar (r = |0.3| see **Fig. 4A**). Prediction quality depended strongly on the imposed delay (**Fig. S8A**). WF-Ca^2+^ occurrences successfully predicted fMRI QPPs with the best correlation between predicted fMRI QPP and original fMRI QPP after applying a delay of 4 frames (equal to 7.2 second delay) to the WF-Ca^2+^ data (**Fig. S8A**). A significant correlation was found for both positive and negative occurrences, between the original and predicted values (r = 0.71/0.70, positive and negative, respectively, p < 0.001, Bonferroni-corrected). The predicted QPP at the optimal delay reproduced the major network-organized structure of the original fMRI QPP (**Fig. S8B**), supporting the idea that WF-Ca^2+^ dynamics precede and predict the corresponding fMRI-QPP with a temporally shifted relationship consistent with a delayed hemodynamic transformation.

## 3. DISCUSSION

Understanding how dynamics in fMRI data reflect neuronal activity is a central challenge in basic and clinical neuroscience research^26,28^. Simultaneous WF-Ca^2+^ and fMRI, enables investigations of resting-state dynamics which unfold across a large FOV using complementary sources of contrast. Here, we derive group- and subject-specific quasi-periodic patterns (QPPs) from simultaneously acquired WF-Ca^2+^ and fMRI data using an established analytical framework^8,10^. The QPPs, derived from each modality, exhibit a high degree of similarity in their spatiotemporal structure. This convergence is not unexpected, as QPPs have been shown to account for a substantial portion of time-averaged, static, functional connectivity, and we observe that static functional connectivity structure is largely preserved across WF-Ca^2+^ and fMRI modalities^29,32,35^. Alignment of WF-Ca^2+^ and fMRI derived QPP occurrences across modalities was maximal at a 3.6-5.4 second delay which matches the expected hemodynamic lag in mice under light anesthesia^7,47^. Further, WF-Ca^2+^ QPP occurrences successfully predicted the spatiotemporal structure of the fMRI-derived QPP, when accounting for a similar temporal delay. Altogether affirming the neuronal origins of spontaneous dynamics in BOLD-fMRI data. Together, our findings support that QPPs in BOLD-fMRI reflect large-scale spatiotemporal dynamics with a neuronal basis, consistent with low-frequency neuronal activity^48^.

Our results extend prior work in rats that has linked QPPs to infraslow neuronal activity using point LFP recordings or global pharmacological manipulation to suppress slow neuronal oscillations (a T-type Ca^2+^ channel blocker)^14,46,49^. LFP recordings are spatially limited and cannot capture dynamics across distributed networks. Suppression of slow waves dramatically reduces the occurrence rate of QPPs (and correlation strength with the fMRI time course) but does not allow for the study of relationships between neuronal activity and the BOLD-fMRI signal. The large FOV afforded by WF-Ca^2+^ imaging provides a measure of spontaneous neuronal activity from the cortical surface (here, comprising 50 ROIs) enabling measures of distributed networks. Recordings of spontaneous neuronal activity alongside the BOLD-fMRI signal allows for regional, and subject-specific differences between complementary sources of contrast to be studied in health, across different developmental stages, and models of injury or disease.

An established and salient feature of QPPs, measured with fMRI in rodents and humans^12,15^, is an oscillation of activity between canonical functional networks^2,4^. In humans, three QPPs are typically reported. The first typically recapitulates the global signal^12,25,50^; the second is typified by an oscillation between the LCN and DMN while the third describes an oscillation (anticorrelation) between SS/MO areas (part of the LCN) with VIS areas. Here, the QPPs, recovered in both imaging modalities after regression of the global signal, share features of both the second and third QPPs that have been descried in the human literature and are in-line with previous work in rodents^15,16^.

Although cross-modality agreement between WF-Ca^2+^ and fMRI derived QPPs was high overall, some differences in their spatiotemporal structure were evident. Specifically, regions within the DMN, the ACA and PL, showed diverging patterns between the WF-Ca^2+^ and fMRI derived QPPs. Sensitivity to anesthesia, differences in vascular architecture, neurovascular coupling, vascular responsiveness, or neuromodulatory tone could all affect the representation of neuronal activity in the BOLD-fMRI signal in a regionally-dependent biologically meaningful manner^51–53^. Consistent with this, recent work combining wide-field voltage imaging and hemodynamics demonstrates that while large-scale activity patterns are shared across modalities, their spatial organization and coupling strength can vary with behavioral state and across cortical regions^48^. Further, the biological basis for neural activity is based on widespread GCaMP expression from a pan-neuronal Synapsin promoter introduced via AAV in neonates. It is possible that calcium transients, buffering and reporter gene expression are different in ACA and PL versus other brain regions, explaining the observed focal differences between modalities. These findings suggest that individual DMN nodes may contribute differently to the generation and expression of QPPs, revealing heterogeneity within the network that is in-line with the literature^54,55^. Understanding these observations will take future study that will rely heavily on multimodal implementations. Importantly, it is not expected that WF-Ca^2+^ and fMRI show a one-to-one correspondence. If they did, their co-implementation would be unnecessary. The QPP derived from WF-Ca^2+^ imaging data showed stronger correlations with the imaging data and more frequent occurrences than the fMRI derived QPP. This was unsurprising given the inter-modality differences in SNR and sources of contrast. In addition, the higher temporal resolution of WF-Ca^2+^ imaging raises the possibility that faster or more transient representations of these dynamics may also be present in the optical data, potentially similar to the motifs described at finer timescales in prior work^56^.

In-line with the data in this study being of high-quality, a QPP resembling the group-derived QPP was recovered from the majority of subjects (using either modality). Differences between subjects, evident at the group- (subject 08) or subject-level (subjects 02, & 06-08) were investigated using measures of data quality (tSNR, motion, and functional connectivity specificity) that are often cited for subject inclusion/exclusion^27,42,57^. Notably, cross-subject, subject-to-group, and cross-modality correspondences were not easily linked to any typical measures of data quality (**Fig. S3-5**). Deviations between the group- and subject-derived QPPs indicated skewed temporal characteristics in the WF-Ca^2+^ imaging data or a lack of a recoverable pattern in the fMRI data (usually, despite strong QPP presence in the simultaneously acquired WF-Ca^2+^ imaging data). Investigating these patterns more thoroughly will require a larger dataset which should include awake as well as anesthetized subjects^58–60^. However, a strength of the multimodal imaging data, and the QPP analysis framework, is the ability to interrogate measures of dynamic brain activity at the single subject level, in-line with investigating individual variability and developing methods for precision medicine. It will of great interest to understand how these multi-modal QPPs vary as a function of arousal in awake mice.

Simultaneous multimodal imaging methods are challenging to implement but necessary for studying the dynamics of spontaneous or resting-state brain activity using complementary sources of contrast. Functional MRI is a modality of critical translational importance given that it can be applied in both human and animal subjects to study large-scale brain activity patterns^27,28^. Yet, the indirect nature of the BOLD-fMRI signal makes a mechanistic interpretation of these data challenging, although recent advances, i.e. Ultrafast fMRI acquisitions, provide improved access to brain dynamics^23,24^. Simultaneous WF-Ca^2+^ and fMRI is an experimental framework for studying spontaneous brain dynamics using complementary sources of contrast. By linking and predicting fMRI-derived QPP structure from neuronal activity, the application of QPP analyses in this work sets the stage for extending to different neural subpopulations (e.g., excitatory- and/or inhibitory- neurons^34^) or other fluorescent signals (e.g., acetylcholine^61,62^) and methods of measuring dynamic brain activity patterns, e.g. the co-activation pattern analyses, which also has demonstrated robust cross species organization of CAPs in fMRI data^2,63^. The findings in this study anchor spontaneous activity patterns measured with BOLD-fMRI signals in concert neuronal activity and establish a translational platform for probing how circuit dynamics change during development and are affected by injury, disease, and treatment.

## 4. METHODS

All procedures were performed following the Yale Institutional Animal Care and Use Committee (IACUC) guidelines and in agreement with the National Institute of Health Guide for the Care and Use of Laboratory Animals. Mice were group housed on a 12-hour light/dark cycle with *ad libitum* access to food and water. C57Bl/6 wild-type mice, N = 8 (4/4 male/female), were used for this study. All mice underwent simultaneous WF-Ca^2+^ and fMRI at 6 months-of-age. The data included here are a subset of a cohort that we have previously published^32,44^. In our previous work, the animals in this study were used to assess differences in static functional connectivity – using the fMRI data only – between wild-type animals and a model of Alzheimer’s disease^44^. The data in this study were also used in a multimodal analysis of awake versus anesthetized mice – again using static functional connectivity^32^. No analyses of dynamic activity have been investigated using these data.

### 4.1 Surgical procedures for WF-Ca^2+^ imaging

#### GCaMP6s Ca^2+^ indicator introduction

We have previously described this procedure^32,41,44^ (**Fig. 1A**). Briefly, on post-natal day 0 (P_0_) pups are anesthetized using ice for 2–3 minutes then transferred to a cool metal plate. A light microscope is used to visualize the transverse sinuses throughout the procedure. Sterilized fine scissors (Fine Science Tools, CA, USA) are used to make two small incisions (∼2 mm) in the skin above each transverse sinus. A glass capillary tube (3.5’ #3-000-203-G/X, Drummond Scientific Co, PA, USA), pulled using a P-97 pipette puller (Sutter Instruments, CA, USA) is used to inject the virus. Pipettes are filled with mineral oil (M3516, Sigma-Aldrich, NY, USA) and attached to a Nanoject III (Drummond Scientific Co) and MP-285 micromanipulator (Sutter Instruments). Most of the mineral oil is ejected and replaced with vector solution, an adeno-associated virus (AAV) for GCaMP6s expression, purchased from https://www.addgene.org/; AAV.Syn.GCaMP6s.WPRE.SV40 - #100843-AAV9. The pipette tip is lowered through the skull and into the lumen of the transverse sinus (300–400 mm below the skull surface). With no delay, 2μL of vector solution is injected at a rate of 20nL per second. The pipette is retracted after a 5 second delay, and the procedure repeated targeting the opposite hemisphere. Delivery is verified by observed blanching of the vessel. With the two injections complete, the skin is sealed with VetBond and the pup is placed on a warming pad. The procedure typically lasts 10 minutes. The total volume of AAV_9_ virus injected per pup is 4μL. To minimize rejection by the mother, a whole litter is removed from the home cage and kept on a warming pad until each pup has undergone the procedure. Once all pups are alert, they are returned to the cage, together, and gently rubbed with bedding to reduce foreign odors. In our hands, this procedure yields a near 100% success rate. Here, all N = 8 injected animals showed expression levels of GCaMP6s sufficient for WF-Ca^2+^ imaging in adulthood (**Fig. S3C**).

#### Head-plate implantation for permanent optical access and head immobilization

We have previously described this procedure^32,44^ (**Fig. 1B,C**). Briefly, all mice were initially anesthetized with isoflurane (5% in 70/30 medical air/O_2_) and head-fixed in a stereotaxic frame (KOPF, USA). Isoflurane was then reduced to 2%. An eye ointment was applied, and meloxicam (2 mg/kg body weight) was administered subcutaneously along with a local injection of bupivacaine (0.1%) at the incision site. The scalp was shaved and washed with betadine and 70% ethanol (three times). The scalp was surgically removed, and the skull cleaned and dried. Antibiotic powder (Neo-Predef) was applied to the incision site and isoflurane reduced to 1.5%. Skull-thinning of the frontal and parietal skull plates was performed using a hand-held drill (tip diameters 1.4mm and 0.7mm). Superglue (Locite) was applied to the exposed skull, followed by transparent dental cement (C&B Metabond). A pre-cut (Neurotechnology Core, USA) all-glass head-plate was attached to the dental cement before solidification. Mice recovered for at least 4 weeks before imaging.

### 4.2 Simultaneous WF-Ca^2+^ and fMRI data acquisition

Multimodal data were collected as we previously described^32^. Data were acquired using an 11.7T MRI scanner (Bruker, Billerica, MA) and our custom-built MRI-compatible WF-Ca^2+^ imaging equipment^29,32,34^.

#### MRI

Data were acquired using ParaVision 6.0.1. Mice were scanned while lightly anesthetized with isoflurane and free breathing (0.5-0.75% in 70/30 medical air/O_2_). Body temperature was monitored (Neoptix fiber), maintained with a circulating water bath (36.6-37°C), and recorded (Spike2, Cambridge Electronic Design Limited). Imaging data and physiological recordings were synchronized using Master-8 A.M.P.I. and Spike2.

#### Structural MRI

For image registration as previously described^29,34^: (1) A whole-brain isotropic 3D image using a multi-spin multi-echo (MSME) imaging sequence, with 0.2×0.2×0.2mm^3^ resolution, repetition time (TR) of 5500ms and echo time (TE) of 20ms, 78 slices, and 2 averages (11 minutes 44 seconds). (2) A fast-low-angle-shot (FLASH) time-of-flight (TOF) angiogram (FOV, 2.0×1.0×2.5cm^3^, covering the cortex), TR/TE of 130/4ms, and resolution of 0.05×0.05×0.05mm^3^ (18 minutes).

#### Functional MRI

Data were acquired using a gradient-echo echo-planar imaging (GE-EPI) sequence: TR/TE of 1.8s/11ms, 334 volumes, 25 slices, and resolution of 0.31×0.31×0.31mm^3^ resulting in near whole-brain coverage (∼10 minutes per run). Three runs were acquired per mouse (total of 30 minutes of fMRI data per mouse).

#### WF-Ca^2+^ imaging

Data were acquired simultaneously with fMRI using CamWare software version 3.17 as previously described^32^. During data acquisition, GCaMP-sensitive (470/24nm) and GCaMP-insensitive (395/25nm) illumination was interleaved at 20Hz using the LLE 7Ch Controller from Lumencor, with a 40ms exposure time per wavelength. To avoid artifacts from the rolling shutter, the imaging sequence included 10ms gaps or blank periods between all 40ms GCaMP-sensitive/-insensitive illumination periods. The raw WF-Ca^2+^ imaging data had a spatial resolution of 25×25μm^2^ and FOV of 1.4×1.4cm².

### 4.3 Data processing

#### Functional MRI

Data processing was performed using RABIES v0.4.8^42^ as we have described previously^32,44^. All fMRI data and isotropic MSME structural images were processed together. Within native subject space, the fMRI timeseries underwent slice time correction and head motion estimation. Structural images were corrected for inhomogeneities using N3 nonparametric nonuniform intensity normalization. A within-sample anatomical template was created by averaging all structural images following non-linear registration. This image was then registered to our in-house template which has been previously shared and registered to the Allen Institute Common Coordinate Framework Reference Atlas (CCfv3)^34^. For each fMRI run, a representative mean image was derived by averaging the motion corrected timeseries. These averaged images were corrected for intensity inhomogeneities before being non-linearly registered to the isotropic structural MSME image acquired from the same mouse. This step minimizes distortions caused by susceptibility artifacts. The fMRI data were then moved to common space using four concatenated transformations: (1) motion correction, (2) mean fMRI to individual MSME, (3) individual MSME to within-dataset template, and (4) within-dataset template to out-of-sample in-house template. Data were resampled to the template resolution (0.2×0.2×0.2mm³). Registration quality was visually inspected using the RABIES quality control report. Timeseries normalization to common space was performed, and the six-parameter motion estimates regressed from the timeseries. Frame scrubbing with a conservative framewise displacement (FD) of 0.075mm threshold was applied. Data were filtered [0.008-0.2Hz], and 30 seconds of data discarded from the beginning and end of each run to avoid filter-related edge artifacts. Cerebrospinal fluid and white matter signals and the global signal were regressed. Data were smoothed with a σ = 0.4mm (**Fig. S1C**).

#### WF-Ca^2+^ imaging

See **Fig. S1C**. For each run, 300 timepoints were discarded from the beginning and end, to match the fMRI time series. Data were smoothed with a σ = 0.1mm and down sampled by a factor of two in each spatial dimension. The GCaMP-insensitive channel was regressed from the GCaMP-sensitive channel, and ΔF/F_0_ (F = fluorescence) computed for each run. After non-linear transformation to common space, the WF-Ca^2+^ imaging data were filtered [0.008-0.2Hz] and temporally down sampled to match the fMRI data. Timepoints corresponding to those censored in the fMRI timeseries (e.g., FD > 0.075mm) were removed from the WF-Ca^2+^ imaging timeseries.

#### Multimodal data registration.

The WF-Ca^2+^ imaging data were registered to the in-house anatomical template using three transformations: (1) WF-Ca^2+^ to MRI angiogram, (2) MRI angiogram to MSME, and (3) MSME to in-house template. These steps were performed using customized tools developed within the BioImage Suite Web (BIS) software package (http://bioimagesuiteweb.github). For more details see **Fig. 2** in ^34^.

### 4.4 Data analyses and statistics

All analyses were performed on MATLAB (v.R2021b). Unless stated otherwise, all analyses adopt Pearson’s correlation (*corr*, MATLAB) with Bonferroni correction. Statistical significance after correction is shown as *p<0.05, **p<0.01 and ***p<0.001.

#### 4.4.1 Data quality metrics

##### Temporal SNR

Functional MRI tSNR was computed as previously described^32^ using RABIES^42^ and [2]. Briefly, *μ_t_* is the mean signal intensity (SI) of a voxel over the course of a run and *σ_t_* is the standard deviation (SD) of the voxel’s SI over the same period. Similarly, the WF-Ca^2+^ imaging tSNR was computed using [2] in MATLAB where *μ_t_* is the mean SI of a pixel within the frame range 1,000-1,500 (50 seconds extracted from the middle of each run).

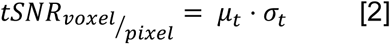

##### Head motion

Motion in the fMRI data was estimated using RABIES^42^, as previously described^32^, using a frame-to-frame displacement threshold (FD) of 0.075 mm. Data excluded based on motion in the fMRI timeseries were also excluded from the WF-Ca^2+^ imaging dataset (**4.3**).

##### Multimodal “specific connectivity”

As previously described^32,57^. Functional connectivity between homotopic SSp (SSp-mouth right/left), SSpR ↔ SSpL, was plotted against connectivity between SSpR and the right ACA, SSpR ↔ ACAR. Data with high connectivity strength (r > 0.1, z-scores) in SSpR ↔ SSpL and concurrent low connectivity strength (r < 0.1, z-scores) in SSpR ↔ ACAR are considered desirable^57^.

These data were plotted alongside a reference distribution created from randomly chosen homotopic node pairs, and inter-hemispheric node pairs (n = 1,000 iteration).

#### 4.4.2 Connectome and timeseries computation

Connectomes were computed using BIS software by averaging fluorescent WF-Ca^2+^ or BOLD-fMRI signals within each ROI, computing inter-regional Pearson’s correlation, and applying a Fisher transformation (**Fig. S1D**). Connectomes were computed for each run using data residing within the in-house template space. For fMRI data, a 172-region whole-brain atlas was used. For WF-Ca^2+^ imaging data, a 60-region cortical 2D atlas was used. Regions were selected from the fMRI 172-region atlas which corresponded to the 60 cortical regions in the atlas applied to the WF-Ca^2+^ imaging data (**Fig. S1**). Both atlases have been reported previously and openly shared through BIS^32,43^. ROI x timepoints timeseries data for QPP analyses were computed in the same way using BIS. Ten ROIs were excluded based on poor coverage (**Fig. S1**) leaving 50 cortical ROIs for all following analyses.

#### 4.4.3 Quasi-periodic pattern analyses

Quasi-periodic pattern analyses were performed as described previously, using the QPPLab toolbox, (see ‘Code Availability’ below)^10–12,14,19^. QPP detection identifies recurring spatiotemporal patterns in large-scale brain activity by iteratively refining a candidate template through sliding-window correlation with the full timeseries. For each imaging modality (WF-Ca^2+^ and fMRI), ROI timeseries data from all mice and runs were concatenated to form a single dataset (preserving the temporal order within runs). Each time point in the dataset representing the activity of 50 cortical regions (**Fig. 2**).

##### QPP template initialization

The algorithm is initialized by randomly selecting a short contiguous segment of the concatenated ROI timeseries to serve as an initial spatiotemporal template. Equation [1] (**Results**) establishes the QPP window length – here, 5.5 frames, thus rounded up to 6.

##### Iterative template refinement

Following initialization, the template is iteratively refined through a sliding-window correlation procedure. At each iteration, the template is slid across the concatenated dataset in a maximally overlapping framework. For every window, the Pearson correlation is computed between the template and data segment of equal length.

This procedure generates a template-correlation time course that reflects the similarity between the candidate pattern and the data at each timepoint. Timepoints exceeding a predefined correlation threshold are considered candidate occurrences of the pattern. In-line with prior work, both positive and negative correlations were considered to capture in-phase and anti-phase presentations of the template. Specifically, correlations r > |0.1| for iterations 1, 2, and 3, and r > |0.2| thereafter (**Results**).

All data segments corresponding to suprathreshold correlations were extracted and averaged to generate a refined template. This refined template was then used in subsequent iterations of the sliding-correlation procedure. Iterations continued until the template stabilized (no changes in QPP).

##### Multiple initializations

To reduce dependence on the initial template, the discovery procedure was repeated using different randomly selected initialization templates, see^10^. The resulting candidate templates were compared, and the refined template with the greatest summed occurrence correlation strength was retained as the final QPP.

##### Detection of QPP occurrences

QPP occurrences within each mouse/run are identified by correlating the QPP with the corresponding ROI timeseries using the same sliding-window correlation procedure as above. Timepoints exceeding r > |0.1| (in-phase and anti-phase) are marked as discrete QPP events. The resulting correlation time course provides a continuous measure of QPP expression within the data, while applying a threshold identifies individual occurrences.

#### 4.4.4 Cross-modal correlation of the QPP

To quantify the spatial correspondence between WF-Ca^2+^ and fMRI derived QPPs, we correlated the vectorized ROIs (at each QPP timepoint) across modalities and displayed a vector of correlation x timepoint (**Fig. 3E**). Then, to recover the spatial component, the WF-Ca^2+^ and fMRI QPPs were z-scored within region. The multimodal QPPs were then multiplied at each timepoint (**Fig. 3F**). Cross-modal QPP similarity was assessed further using the z-scored values by computing the Pearson correlation between the WF-Ca^2+^ and fMRI QPPs across ROIs within timepoint (similar to **Fig. 3E**, but after z-scoring). Statistical significance was evaluated using a permutation test in which ROIs were randomly shuffled (2,000 iterations) to generate a null distribution of correlation values. Resulting p-values were corrected for multiple comparisons across frames using Bonferroni correction and displayed as a histogram: null in grey observed values in red (**Fig. 3G**).

#### 4.4.5 Cross-modal temporal alignment of WF-Ca^2+^ and fMRI QPPs

Cross-correlation was performed on WF-Ca^2+^ and fMRI derived QPPs. To recover the expression of the group QPP within the multimodal data from each mouse/run, we averaged across the identified occurrences (r = |0.1|). Within run, the averaged WF-Ca^2+^ and fMRI QPP timeseries was normalized and cross-correlated (*xcorr*, MATLAB) using Pearson correlation with and without introducing a temporal lag between modalities. Cross-correlation results were evaluated over a lag window of ± 20 seconds (sampling interval TR = 1.8 seconds) (**Fig. 4C,D**).

Correlations were restricted to lags between 0 and 9 seconds (**Fig. 4D**). For each run, the lag producing the maximum correlation within this interval was identified and reported as the best lag together with its corresponding peak correlation. To test whether correlation strength varied systematically across lags, a repeated-measures analysis of variance (ANOVA) was performed with lag as a within-run factor. Post hoc comparisons across lag values were conducted with Bonferroni correction for multiple comparisons. To visualize the lag-dependent correlation structure, the run-by-lag correlation matrix was displayed as a heatmap. Statistical significance at each lag was evaluated using a one-sample two-sided t-test across runs, followed by Bonferroni correction across lags. Significant lags are indicated on the heatmap (**Fig. 4E**).

#### 4.4.6 Cross-modal QPP switching

Similar to previously work^19^, a ‘QPP swap’ approach was investigated. QPPs derived from one modality were applied to the data of the second modality to generate a ‘QPP swap’ derived occurrence time course. This procedure resulted in four occurrence time courses (two “derived” and two “original”). At each occurrence in the original version, the corresponding 6 timepoints (dictated by QPP half-cycle, in-phase or anti-phase, length) were extracted from both the original and derived occurrence time courses and correlated using Pearson correlation. Thus, for each occurrence in the original occurrence time course, a correlation between the original and ‘QPP-swap’ derived version was obtained.

##### Null model and statistical testing

Statistical significance was assessed using permutation-based null models that disrupted temporal alignment while preserving within-run structure. For each iteration, the original occurrence time course was shuffled within run, and correlations with the ‘QPP-swap’ derived occurrence time course recomputed (n = 2000 iterations). Empirical null distributions were generated across iterations, and two-sided empirical p-values were calculated by comparing observed correlations with the corresponding null distributions. Bonferroni correction applied.

### 4.5 Inclusion criteria

Frames where head motion, FD > 0.075mm (measured using fMRI data), were censored from both imaging modalities. Ten regions at the edge of the cortical surface were excluded due to partial coverage in the WF-Ca^2+^ imaging FOV (**Fig. S1B**). No runs or mice were excluded due to poor registration, low tSNR, or an excessive number of frames being censored due to head motion.

## Acknowledgments

Drs. Joel Greenwood, Paul Shamble, and Omer Mano from Neurotechnology Core of the Kavli Institute for Neuroscience at Yale and the team members of the Magnetic Resonance Research Center at Yale University for their technological expertise. Dr. Joanes Grandjean for critical discussion. Dr. Yonghyun Ha for hardware support. Dr. Ali S. Hamodi and Dr. Michael C. Crair for experimental expertise.

## Author contributions

Conceptualization: FM, EMRL, NX, SK.

Methodology: FM, XS, CH, SK, NX, XP.

Investigation: FM, CH.

Visualization: FM.

Funding acquisition: EMRL.

Project administration: FM, EMRL.

Supervision: EMRL.

Writing—original draft: FM.

Writing—review & editing: all co-authors.

## Data availability

Data are not available in a public repository due to ongoing work by the authors on this dataset. Access can be obtained by contacting the corresponding authors.

## Code availability

Software packages for data processing are available: multimodal data registration and processing of human data (BioImage Suite, https://bioimagesuiteweb.github.io/webapp/), and fMRI preprocessing (RABIES, https://github.com/CoBrALab/RABIES). Code for quasi-periodic pattern analyses can be found here: https://github.com/GT-EmoryMINDlab/QPPLab.

