## Supplementary for "Harmonizing brain rhythms: cortex-wide neuronal dynamics underpin quasi-periodic patterns in resting-state fMRI"

### Supplementary Material

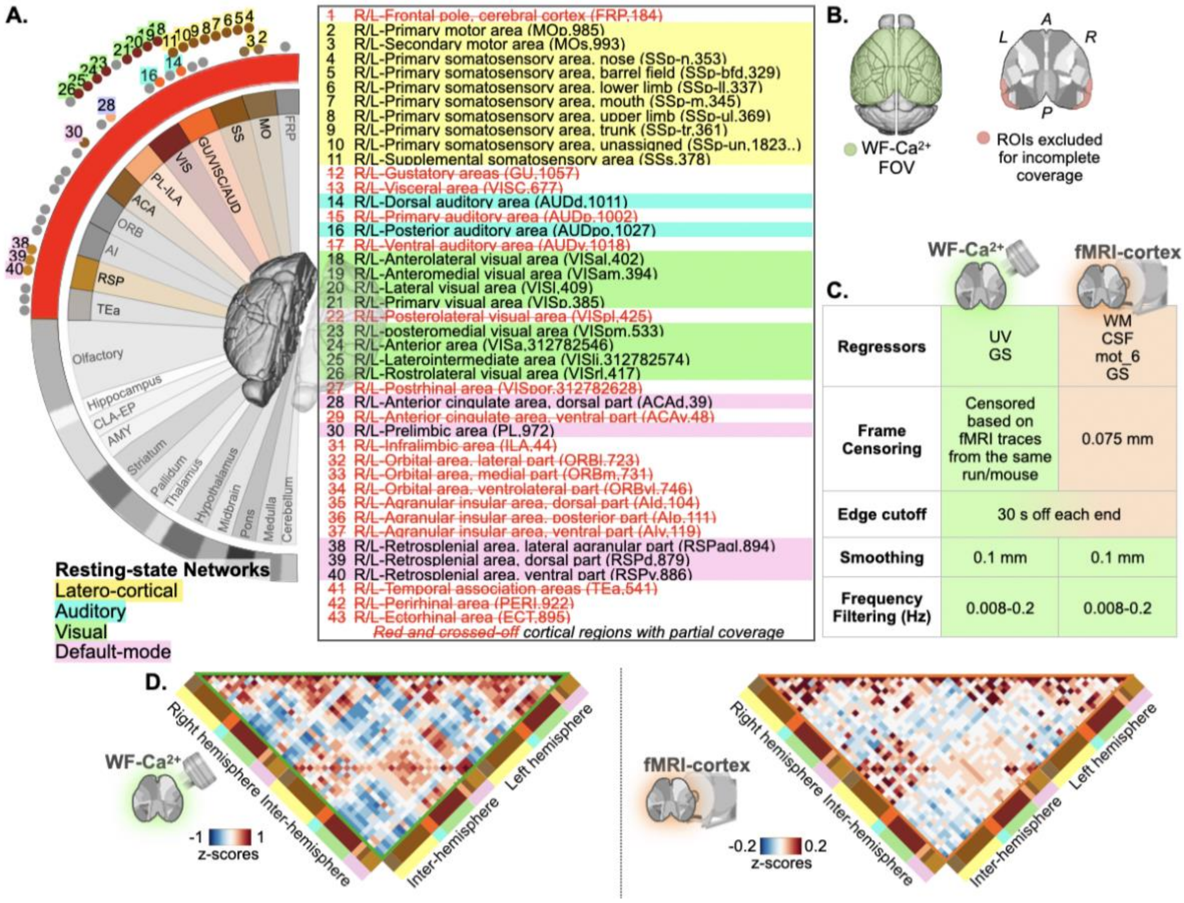


**Fig. S1. Allen Atlas derived brain regions and preprocessing steps. A&B.** Brain regions visible in the WF-$Ca^{2+}$ imaging field-of-view (FOV) used in all analyses. All 50 regions were subdivided into four *a priori* functional networks (**A**, bottom left)**.** Red strike-through regions are not included in the analyses (**A**, right). **C.** Details of preprocessing steps for both WF-$Ca^{2+}$ and fMRI. **D.** Average connectomes for static functional connectivity (z-scored) for . Stereotypical patterns including high FC between bilateral brain regions were observed. UV: ultra-violet (405nm, calcium insensitive background channel); GS: global signal, WM: white matter; CSF: cerebrospinal fluid; mot_6: 6 motion parameters; ROIs: regions-of-interest.

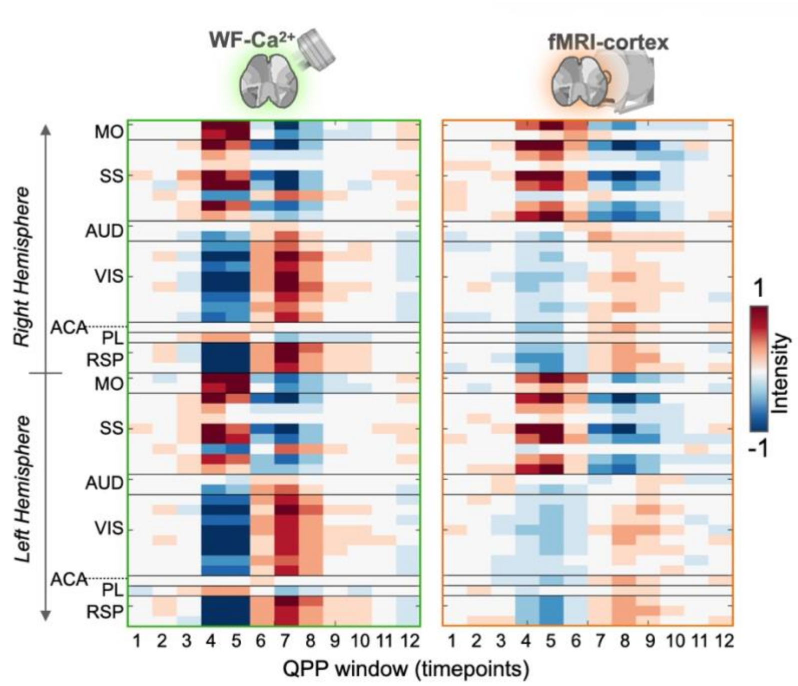


**Fig. S2. WF-Ca$^{2+}$ and fMRI derived QPPs show high bilateral symmetry.** Group QPPs derived from simultaneously acquired WF-Ca$^{2+}$ (left) and fMRI data (right). ROIs are grouped by hemisphere to emphasize bilateral symmetry. ROIs: regions-of-interest, MO: motor, SS: somatosensory, AUD: auditory, VIS: visual, ACA: anterior cingulate, PL: prelimbic, RSP: retrosplenial.

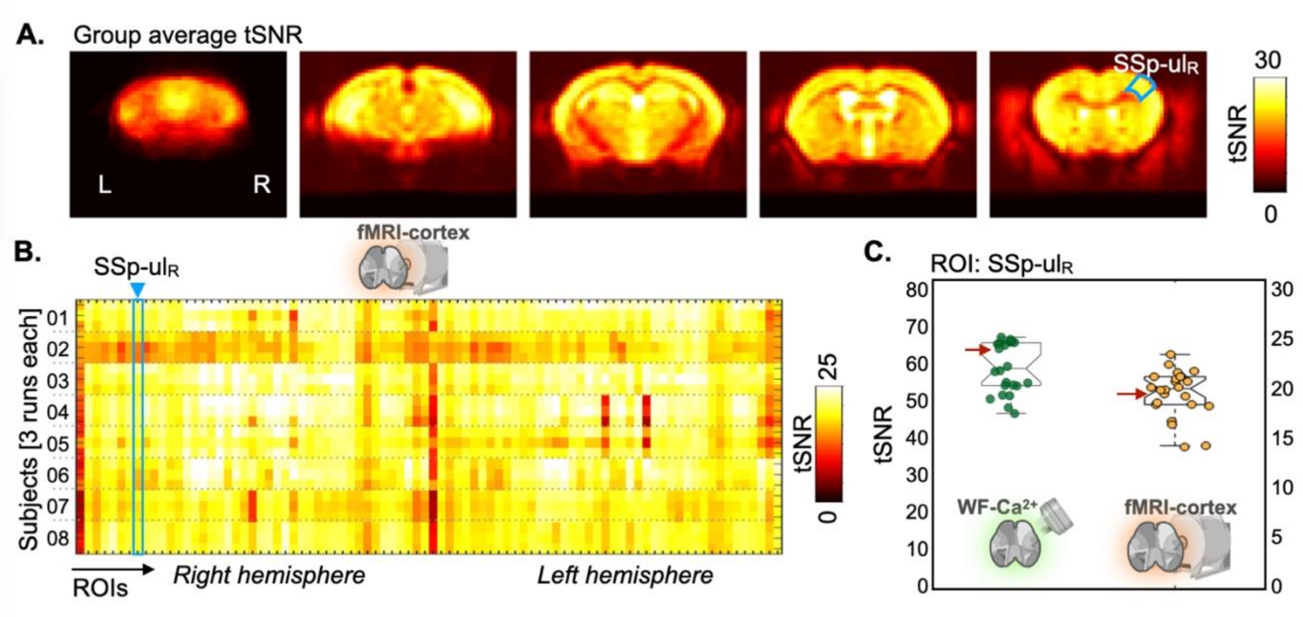


**Fig. S3. Temporal SNR across multimodal measures. A.** Example slices from the group averaged tSNR map generated from fMRI data using RABIES. Indicated in cyan the primary somatosensory, upper limb, region ($SSp\text{-}ul_R$), used as reference for ROI-specific tSNR analyses. **B.** Carpet plots from RABIES which show tSNR estimations for each brain region and mouse/run. In cyan outline and indicated with a reversed triangle, $SSp\text{-}ul_R$ reference ROI. **C.** tSNR across modalities shown for a representative brain region ($SSp\text{-}ul_R$) for WF-$Ca^{2+}$ (left) and fMRI (right). Subject 08 indicated with red arrows. tSNR: temporal signal to noise ratio; L/R: left/right.

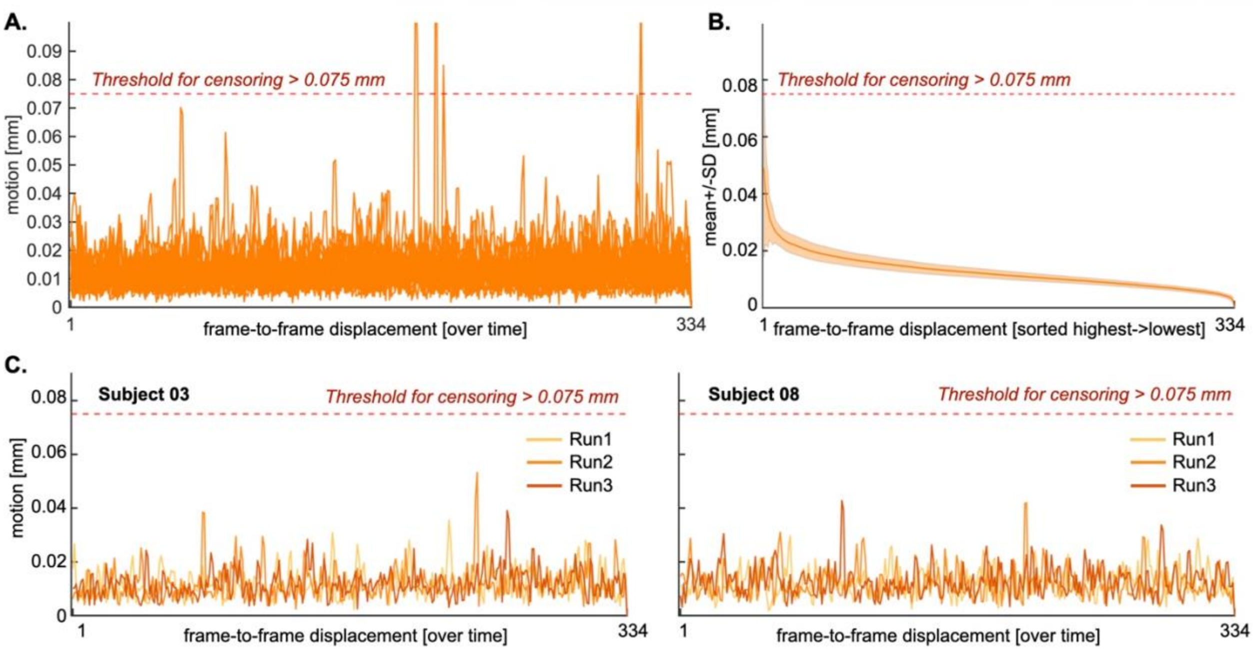


**Fig. S4: Head motion estimates from fMRI data**. **A.** Head motion (frame-to-frame displacement, FD) from each mouse/run generated from fMRI data (RABIES). Frame number (from the beginning to the end of the scan) is plotted on the x-axis. The threshold for frame censoring (>0.075 mm) is indicated on the y-axis. **B.** Mean ± SD head motion (FD) sorted from high-to-low. Frame number is on the x-axis (sorter from high to low). **C&D.** Example of head motion (FD) estimates for subjects 03 and 08, respectively.

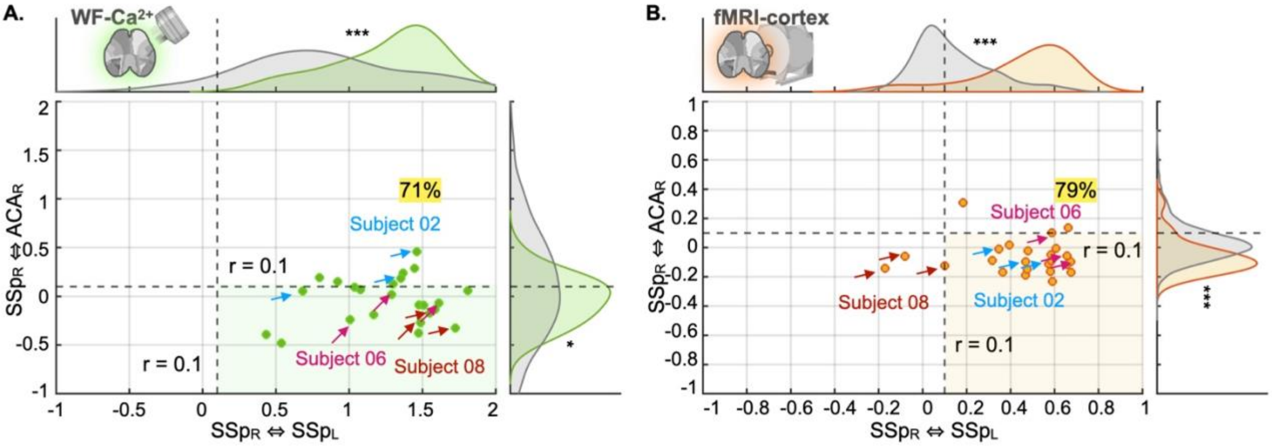


**Fig. S5. Multimodal "specific connectivity".** Scatterplots showing "specific" functional connectivity using WF-$Ca^{2+}$ imaging data (**A**) and fMRI data (**B**). In the "specific connectivity" framework, high ($r > 0.1$, z-scored) $SSp_R \Leftrightarrow SSp_L$ (x-axis) versus low ($r < 0.1$, z-scores) $SSp_R \Leftrightarrow ACA_R$ (y-axis) connectivity strength should isolate 'high-quality' datasets in the lower right quadrant (highlighted in light green/light orange for WF-$Ca^{2+}$ and fMRI data, respectively). Above and to the right of the scatterplots, density distributions display kernel-smoothed observed (solid lines) and reference (dashed lines) distributions. Here, the reference distributions were generated from randomly selected intra- and inter-hemispheric node pairings, as described in [30]. Example subjects 02, 06 and 08 are indicated (cyan, pink, red respectively). Corrected p-values displayed: *$p<0.05$, **$p< 0.01$, ***$p<0.001$.

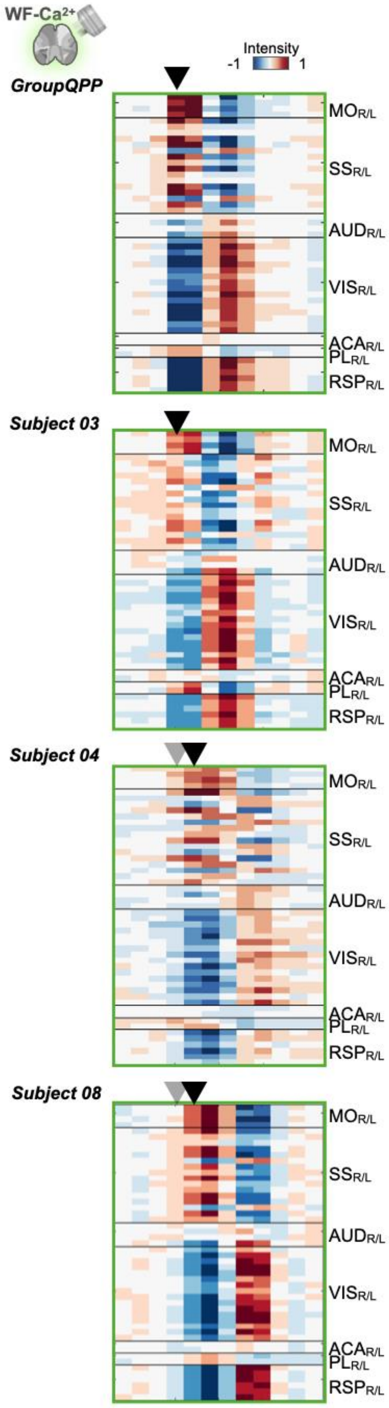


**Fig. S6. Single-subject QPP.** Top: group-derived WF-$Ca^{2+}$, reproduction from **Fig. 3A**, left. Below: example of single-subject QPP derived from within-subject QPP detection. Actual onset of QPP marked with black triangle, and group-reference onset in grey in shifted QPPs.

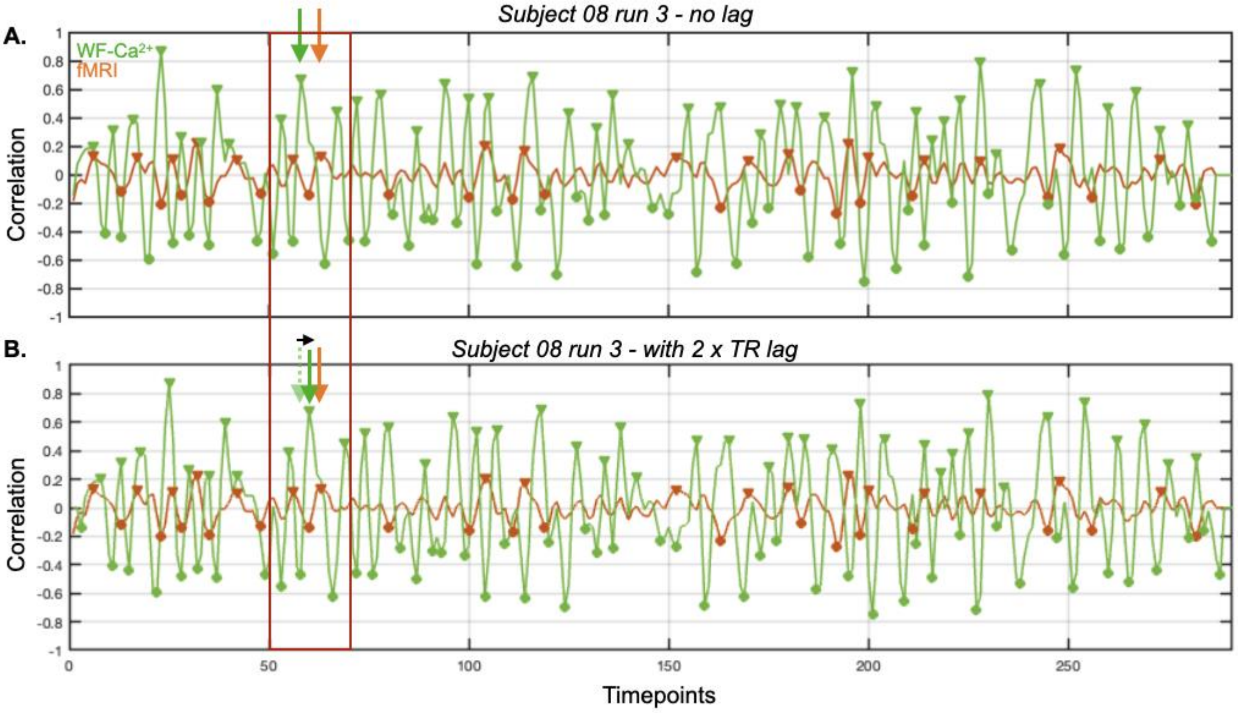


**Fig. S7. QPP occurrences in subject 08 with and without a lag.** Representative multimodal QPP occurrence time courses for subject 08 without and with a temporal lag of 2TRs (**A&B**, respectively). WF-$Ca^{2+}$ imaging occurrence time courses are shown in green. Functional MRI occurrence time courses are shown in orange.

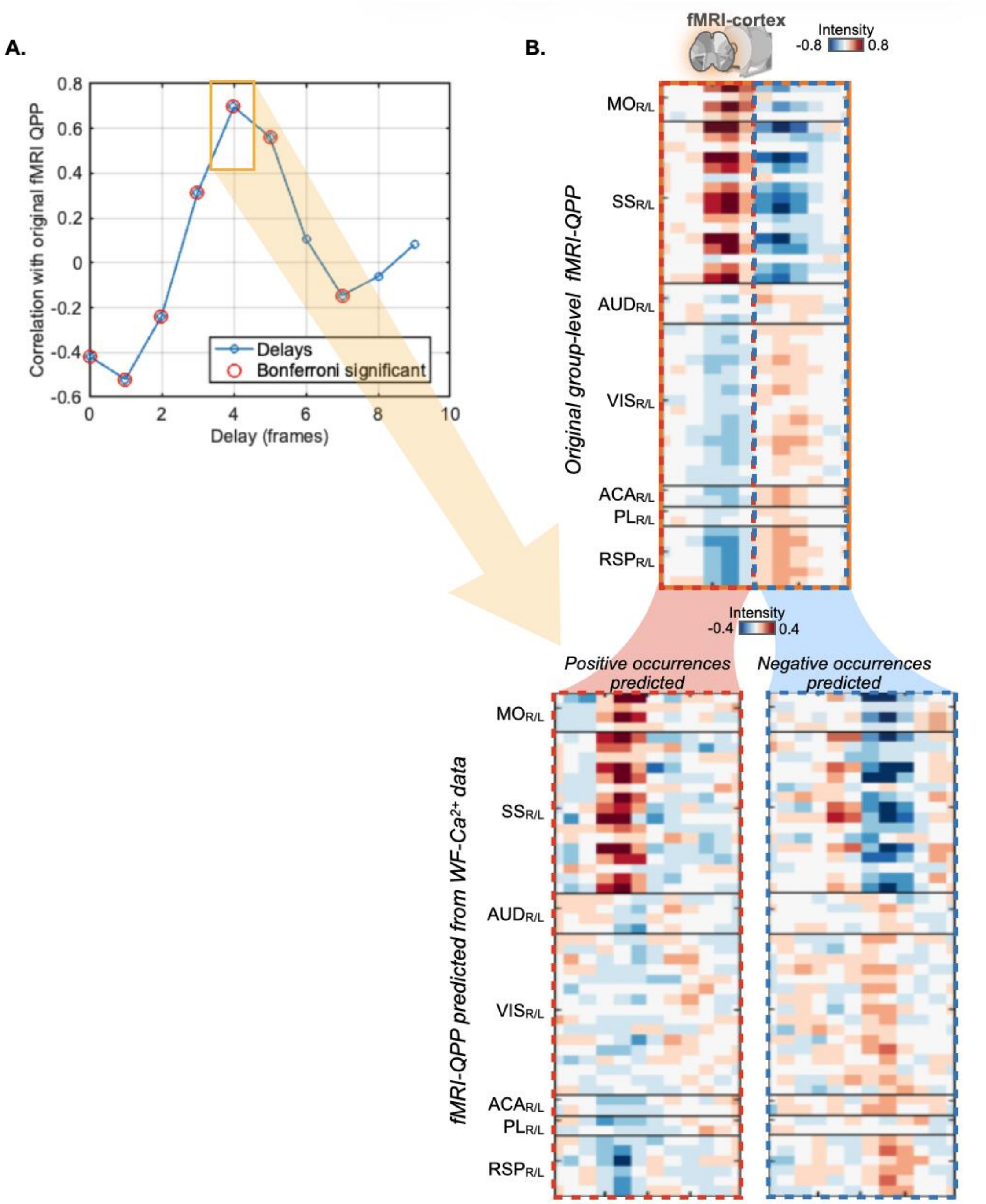


**Fig. S8. WF-Ca$^{2+}$-derived prediction of fMRI QPP structure across phases. A.** Correlation between 'predicted' fMRI-QPP (predicted from WF-Ca$^{2+}$ temporal occurrences) the original fMRI-QPP, plotted as a function of temporal delay *k* (*k* = 0-9 frames). Each point represents the correlation at a given delay, markers indicating delays that survive Bonferroni correction. **B.** Top: Original group-level fMRI QPP showing the canonical spatiotemporal pattern across cortical networks (reproduced from **Fig. 3A**, right). Bottom: fMRI QPPs predicted from WF-Ca$^{2+}$ QPP occurrence times during the mismatched correlation threshold of |0.3|, using positive (left, red dashed line) and negative (right, blue dashed line) WF-Ca$^{2+}$ events separately for optimal time lag (*k* = 4 frames). The WF-Ca$^{2+}$-derived templates recapitulate the biphasic structure of the fMRI QPP, with positive occurrences predicting the main QPP in-phase (left, red dashed lines) and negative occurrences predicting the anti-phase (right, blue dashed lines).
